# PSP-0119: Targeted IRAK4 Degradation as a Novel Therapeutic Strategy for FLT3-Mutant AML

**DOI:** 10.1101/2025.09.30.679569

**Authors:** Negar Khazan, Cameron WA Snyder, Chandhana Ravi, Elizabeth Lamere, Niloy A. Singh, Manoj K. Khera, Jane Liesveld, Srinivasan Ekambaram, Nikolay V. Dokholyan, Myla Strawderman, Kyu Kwang Kim, Rachael Rowswell-Turner, Michael W. Becker, Richard G. Moore, Rakesh K. Singh

## Abstract

Acute Myeloid Leukemia (AML) is a life-threatening hematologic malignancy. Despite recent therapeutic advances, rising incidence rates emphasize the urgent need for identification of new targets and therapies. Roles of interleukin receptor-associated kinases IRAK1/4 are emerging in hematologic and solid malignancies. In AML, IRAK4 mRNA is overexpressed at diagnosis, relapses, in residual disease, and in FLT3-ITD-mutant cells, MDS, MPN, and MDS/MPN-negative subtypes. Compared with hematopoietic stem cells, IRAK4 is elevated in t(15;17), inv(16)/t(16;16), and t(11q23)/MLL subtypes, correlating with poor survival. Here, we disclose anti-AML activity of PSP-0119, a novel IRAK4 PROTAC degrader. PSP-0119, inhibited IRAK4 kinase activity, NF-κβ activity, and IL-1β-induced IRAK4 phosphorylation. In-silico docking revealed interactions in CRBN/IRAK4/PSP-0119 ternary complex. PSP-0119 degraded IRAK4 in FLT3-mutant AML cell lines sparing FLT3-wild-type AML cells, FLT3-wild-type patient samples, and normal bone-marrow. Bulk-seq of PSP-0119 treated MOLM-13 cells revealed downregulation of eNOS, a poor AML prognosticator. PSP-0119 suppressed colony formation, cell viability, and MOLM-13 xenograft growth, and synergized with IRAK1 covalent inhibitor JH-X-119-01. PSP-0119 is metabolically stable, retaining 71% of parent compound at 60 minutes in human liver microsomes. In summary, IRAK4 degradation via PSP-0119 as a promising therapeutic strategy for treatment of FLT3-mutant AML.

## Introduction

Acute myeloid leukemia (AML) is a difficult-to-treat, life-threatening hematological malignancy^1^. Patients with AML face dismal 5-year survival rates, limited to only 10–15% in those aged ≥60 years and 30–35% in younger patients^2^. Patients with relapsed AML face the greatest risk as the current treatments have plateaued in ability to control disease^3-4^. There is an unmet need to identify novel, drug-responsive targets and therapies that directly address the key drivers of AML^5-6^. Myelodysplastic syndromes (MDS) and AML are increasingly linked to dysregulated innate immune and inflammatory IL-1/TLR signaling pathways^7-10^. This reprogramming disrupts multiple downstream mechanisms controlling cell survival, immune dysfunction, chemoresistance, and recurrence by hijacking oncogenic NF-κβ pathways through MyD88 and the interleukin-1 receptor–associated kinases IRAK1 and IRAK4^11^. Recurrent genetic mutations observed in AML and MDS, such as spliceosomes^12^ and epigenetic mutations^13^, have been linked to alterations in IL-1/TLR signaling. Furthermore, published studies^14^, including our ongoing ones^15^, have demonstrated overexpression of key components of IL-1/TLR signaling, including the IL-1 ligand, receptor IL1R and co-receptors in AML and MDS patients. No therapy targeting the IL-1/IL1R/TLR axis has yet been approved for AML, its precursor MDS, or solid tumors^16-17^. The IRAK4 inhibitor CA-4948 (Emavusertib) is currently in clinical trials for myeloid malignancies (NCT04278768); however, because CA-4948 is a tri-kinase inhibitor (IRAK4, FLT3, and CLK), its effects can be confounded by simultaneous targeting of multiple kinases, leaving the impact of selective IRAK4 inhibition untested in AML and MDS models.

Previously, we showed that targeting IL1/IL1R/TLR signaling may orchestrate underlying changes in functionally defined LSC populations observed in AML patients at time of diagnosis and relapse^15^. We also showed that the loss of IL-1β signaling, factors IL1RAP and IL1R1 affect LSC functions *in-vitro* and *in-vivo*, each of which provide rationale to target IL1β/IL1R1 driven signaling in AML. Following which, we developed UR241-2, a novel small molecule compound targeting IRAK4 to counteract IL1/IL1R1 driven signaling in AML. UR241-2 suppresses IL-1β/IRAK1/4 downstream signaling, including the activation of NF-κβ and the phosphorylation of p65 and p38, consequently impeding LSC clonogenicity and engraftment. However, IRAK1 promotes metastasis, migration, invasion, EMT transition and can drive resistance including against IRAK4 in AML and other malignancies^18-19^. IRAK4 carries scaffolding functions in addition to kinase functionality integral to myddosome functions. Scaffolding functions of IRAK4 allow it to assemble protein complexes, influence substrate specificity, and regulate signaling pathways independently of its kinase activity. Hence, we developed PSP-0119^20^, a PROTAC (Proteolytic Targeting Chimers) that can degrade both IRAK1 and IRAK4 to maintain sustained control over AML. PROTAC PSP-0119 could degrade the entire target protein IRAK4, blocking both catalytic and non-catalytic/scaffolding functions, which UR241-2 was unable to achieve.

In this study, we demonstrate how PSP-0119 degrades IRAK4 selectively in FLT-3 mutant AML sparing wild-type AML cells, bone marrow samples from patients with AML, normal human bone marrow samples, and impacts the cells viability, colony formation CFU and xenograft tumor growth. PSP-0119 which has shown notable human liver microsomal stability will act catalytically so that one molecule can eliminate numerous aberrantly active IRAK4 units, providing longer-lasting control over AML and avoiding the risk of resistance development from kinase domain mutations that IRAK4 kinase inhibitors may face.

## Results

Our study shows that IRAK4 is significantly correlated with initial diagnoses, residual, relapse, MDS (myelodysplastic syndrome) or Myeloproliferative neoplasms (MPNs) disorders and FLT-3-ITD (internal tandem duplication) mutation (Figure-1: A-B). MDS and MPN are often the precursors to AML while MDS represents inefficient production of white cells by bone marrow (BM). MPNs are chronic blood cancers characterized by the overproduction of blood cells. On the other hand, FLT3-ITD-mutation is a major driver of AML. FLT3-ITD, are found in ∼30% of newly diagnosed AML cases and can drive disease progression and relapse. Among the AML karyotypes, while IRAK4 is significantly overexpressed in T(11q23)/MLL Inv(16)/t(16:16), and T(15:17) AML phenotypes (Figure-1C), IRAK1 overexpression is significantly associated with AML complex and T(11q23)/MLL (Figure-1E). Kaplan-Meier analysis showed that both IRAK1 and -4 overexpression showed greater risk of mortality upon upregulation (Figure-1D and Figure-1F).

**Figure-1.**
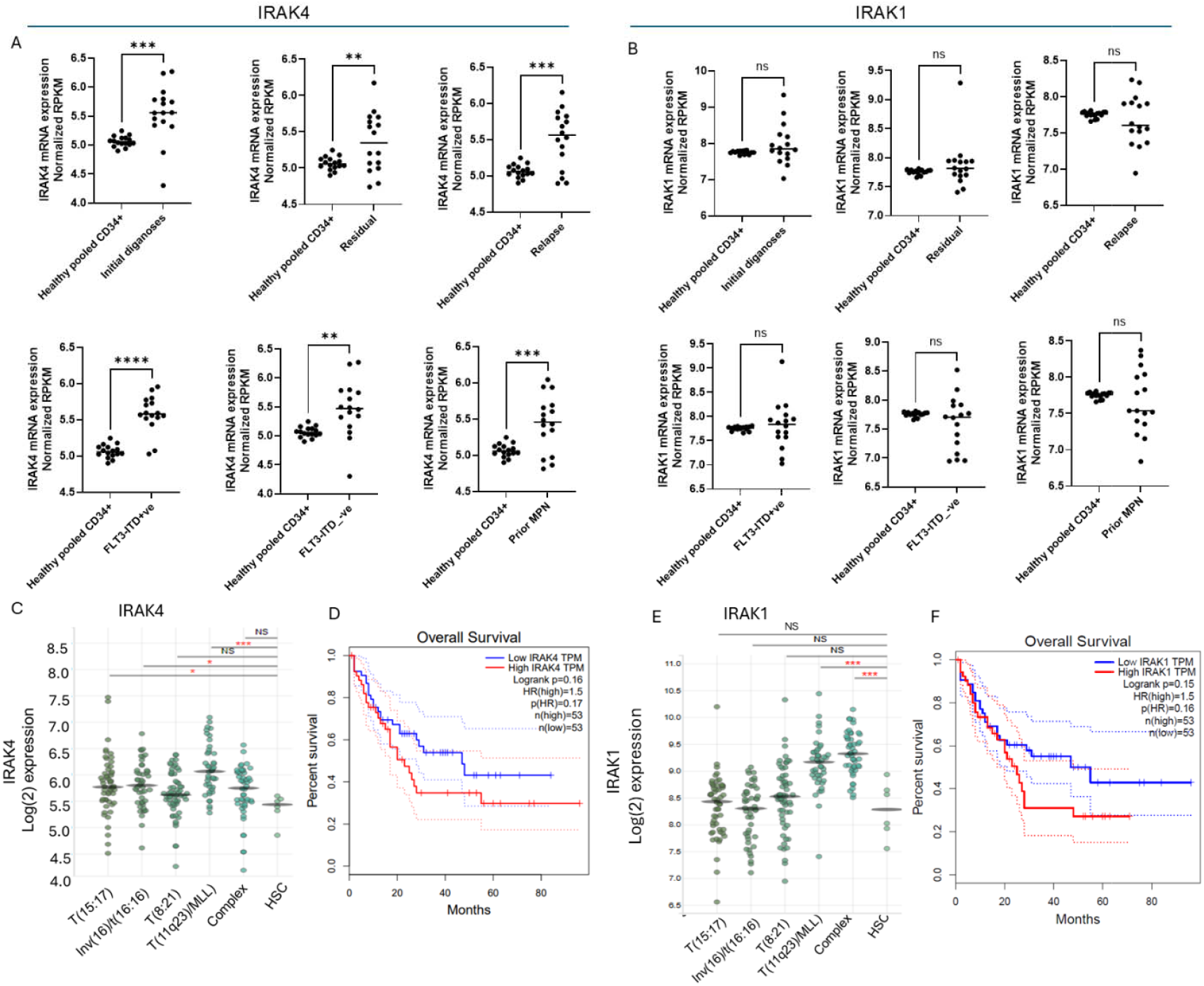
**(A**): Analysis of the AML patient’s microarray database (source: http://vizome.org/aml2/expression_strat/ showed that IRAK4 mRNA is significantly overexpressed in initial diagnosis (p<0.0001, n=16 each), relapse (p<0.001, n=16 each), residual disease (p=0.031, n=16 each), residual (p=0.034, n=16 each), FLT-3-ITD positive (p<0.0001, n=16 each), prior MDS (p=0.03, n=16 each), prior MPN (p<0.0001, n=16 each) and MDS- MPN+ve patients (p<0.0001, n=16 each) compared to healthy pooled CD34+ cells. On the other hand, **(B**): IRAK1 mRNA expression was not significantly upregulated among these AML conditions than healthy pooled CD34+ cells. (**C**): IRAK4 mRNA is significantly upregulated in T(15:17), Inv(16)/t(16:16) and T(11q23)/MLL AML patients (source: Bloodspot.org). (**D**): Kaplan-Meier analysis showed that IRAK4 mRNA overexpressor patients face early and greater proportions of deaths compared to low expressor patients (Source: GEPIA, http://gepia.cancer-pku.cn/detail.php?gene=IRAK4&clicktag=survival). (**E**): IRAK1 mRNA is significantly upregulated in complex and T(11q23)/MLL karyotype presenting AML patients (source: Bloodspot.org). (**F**): Kaplan-Meier analysis showed that IRAK1-mRNA overexpressor patients face early and great proportions of deaths than low expressor patients (Source: GEPIA, http://gepia.cancer-pku.cn/detail.php?gene=IRAK1&clicktag=survival).

To disrupt the IL1/TLR/IRAK4-driven malignant niche in AML, we recently reported a novel IRAK4 kinase inhibitor UR241-2, which suppressed IL-1–induced IRAK1/4 signaling, NF-κβ activation, and phosphorylation of p65 and p38. UR241-2 selectively inhibited leukemia stem cell clonogenicity in primary AML cells at diagnosis and relapses, while sparing normal hematopoietic progenitors, and reduced AML engraftment in leukemic mice^15^. However, targeting IRAK4 remains challenging due to the independence of scaffolding functions from its kinase activity, which ATP-competitive inhibitors like UR241-2 cannot block. PROTACs have emerged as promising therapeutics capable of degrading proteins by combining two binding elements, one targeting the protein of interest and the other recruiting an E3 ubiquitin ligase-linked together. This enables the ubiquitin-proteasome system to degrade the target, eliminating both catalytic and scaffolding functions. PROTACs have demonstrated superior anti-important activity against targets resistant to conventional inhibitors (e.g., alisertib), underscoring the importance of scaffolding functions in tumorigenesis. Designing an effective PROTAC is complex, as the two binding moieties and linker must be synergistically optimized for efficient protein degradation.

To synthesize PROTACs targeting IRAK1 and IRAK4, we selected the pyrazolo-pyridine scaffold of UR241-2 or JH-I-25 to engage the IRAK1/4 moiety within the ternary complexes of IRAK1/CRBN or IRAK4/CRBN. Derivatives of this scaffold consistently exhibit nanomolar inhibition of both IRAK1 and IRAK4. Additionally, the PROTAC construction was designed to occur in the solvent-accessible region, which was expected not to impair IRAK1/4 engagement. Thus, two PROTACs PSP-0102 and PSP-0119 were synthesized with varied CRBN ligand and linker structures (Schema 1-2) (NMR, Mass, HPLC data: Supplementary Figure-S1). PSP-0119 constitutes lenalidomide (CRBN hook) linked with a piperazinyl chain (linker) and UR241-2 (IRAK4 hook) whereas Hotspot kinase assay verified that despite drastic modifications on UR241-2 structure, the kinase inhibition activity of PSP-0119 was not lost and showed IRAK4 kinase inhibition at 2.83nM, comparable to UR241-2 in the hotspot assay (1.81nM). PSP-0119 was about 3-fold more potent than PSP-0102 analog in terms of IRAK4 kinase inhibition (Figure-3A). This study, therefore, focuses on PSP-0119. In silico docking showed that phthalidomide analog portion of PSP-0119 engaged with CRBN and pyrazolo-pyridine portion of the molecule docked to IRAK4 kinase (Figure-3:left).

**Scheme-1.**
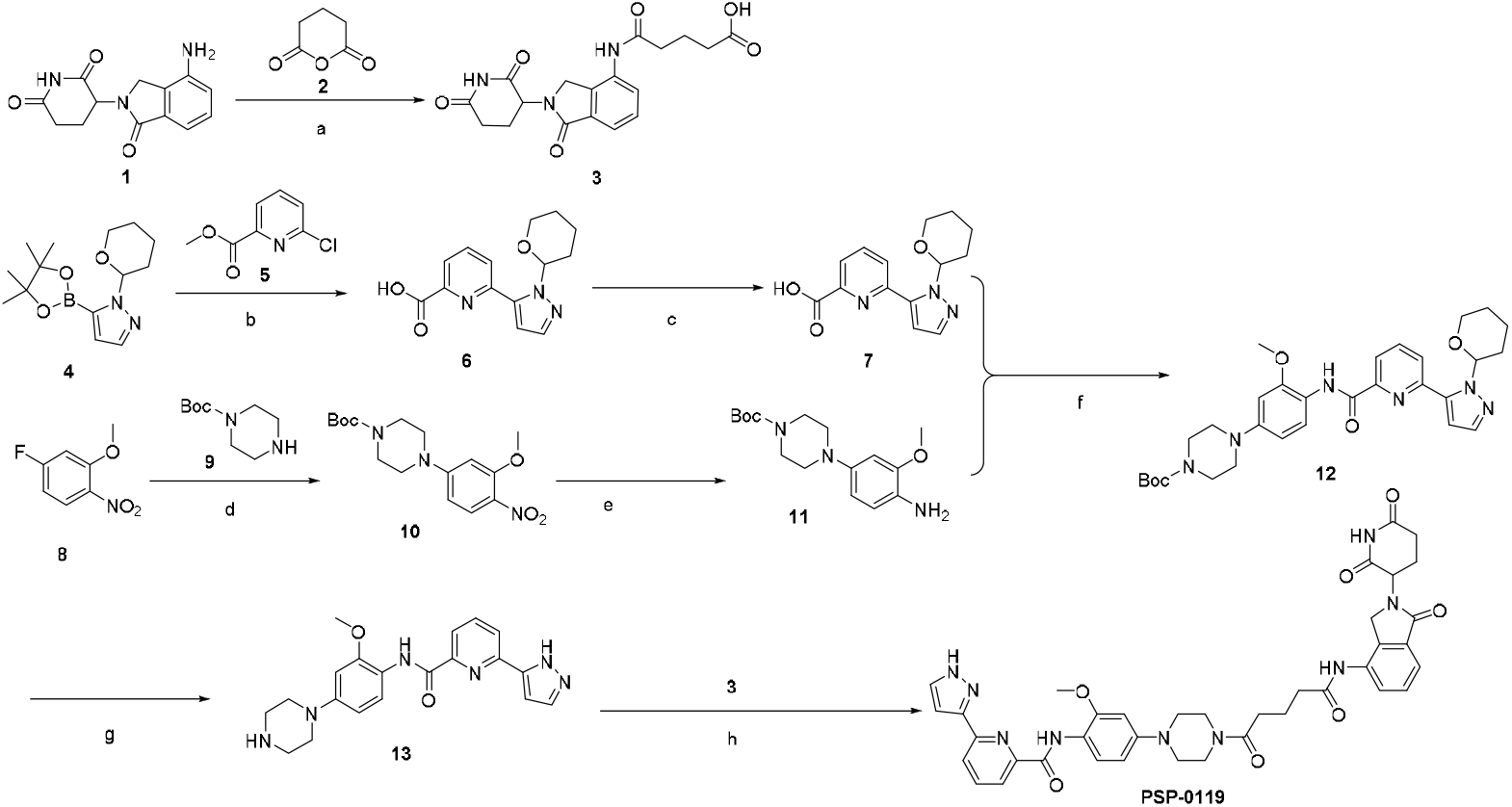
Reagents and conditions: (a) Toluene, reflux, 6 h, 80%; (b) Pd(PPh_3_)_4_, K_2_CO_3_, Dioxane :H_2_O, 80 °C, 16 h, 89%; (c) LiOH.H_2_O, THF: H_2_O, rt, 6 h, 83%; (d) K_2_CO_3_, DMF, 80 °C, 16 h, 88%; (e) H_2_, Pd/C (10%), MeOH, rt, 95%; (f) EDC.HCl, HOBt, DMF, rt, 3 h, 74%; (g) TFA, DCM, rt, 99%; (h) EDC.HCl, HOBt, DMF, rt, 3 h, 64%.

**Scheme-2.**
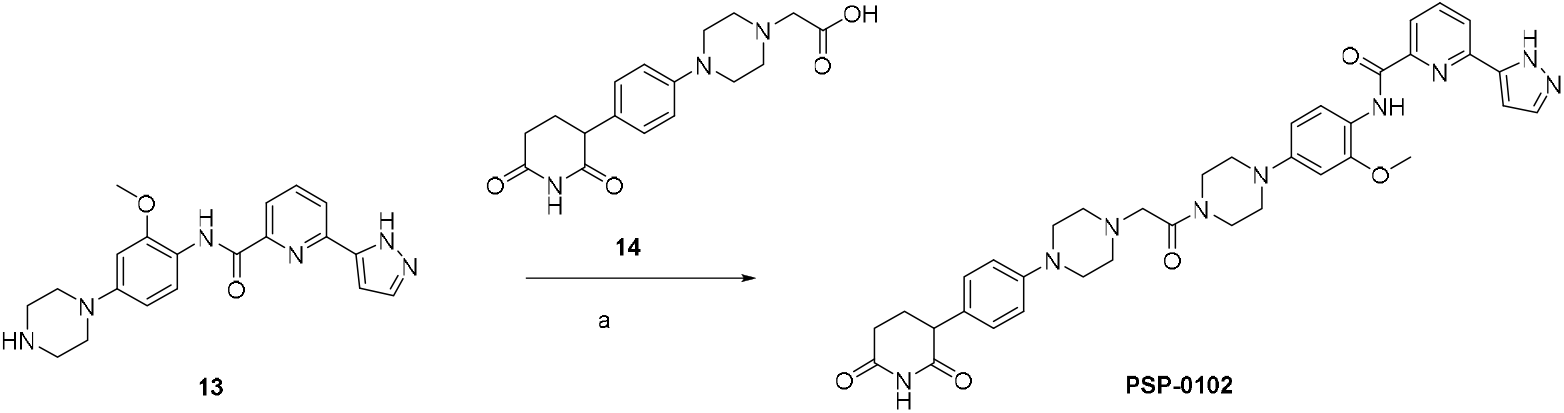
(a) EDC.HCl, HOBt, DMF, rt, 3 h, 70%.

Upon activation, IRAK4 initiates a signaling cascade that activates nuclear factor-kappa β (NF-κβ), driving the expression of pro-inflammatory cytokines and immune response genes. PSP-0119 inhibited NF-kβ activity in THP-1 cells stably expressing NF-kβ-Luc reporter (Figure-2B). PSP-0119 robustly blocked IL1β induced activation (phosphorylation) of IRAK4 in MOLM-13 cells (Figure-2C). We utilized high-resolution X-ray crystal structures of IRAK4 (PDB ID: 8W3X) and CRBN (PDB ID: 6BOY) as starting conformations for our in-silico docking analysis. Protein-protein docking was performed using the HDOCK program, which employs a hierarchical docking algorithm incorporating both template-based modeling and ab initio docking approaches, to generate the IRAK4-CRBN complex. The resulting optimized complex served as the receptor for molecular docking studies of using the Medusa-Dock program^21-23^. The docking focused on the binding pocket defined by the co-crystallized ligand in the IRAK4 structure, ensuring biological relevance of the predicted binding modes. PSP-0119 underwent 100 independent docking iterations to ensure unbiased, thorough sampling of possible binding conformations and to enhance the reliability of the results. PSP-0119 demonstrated superior binding characteristics with the ternary complex, exhibiting a binding energy of -86.5 kcal/mol. These computational findings strongly indicate that PSP-0119 forms a highly stable ternary complex. (Figure-3).

**Figure-2.**
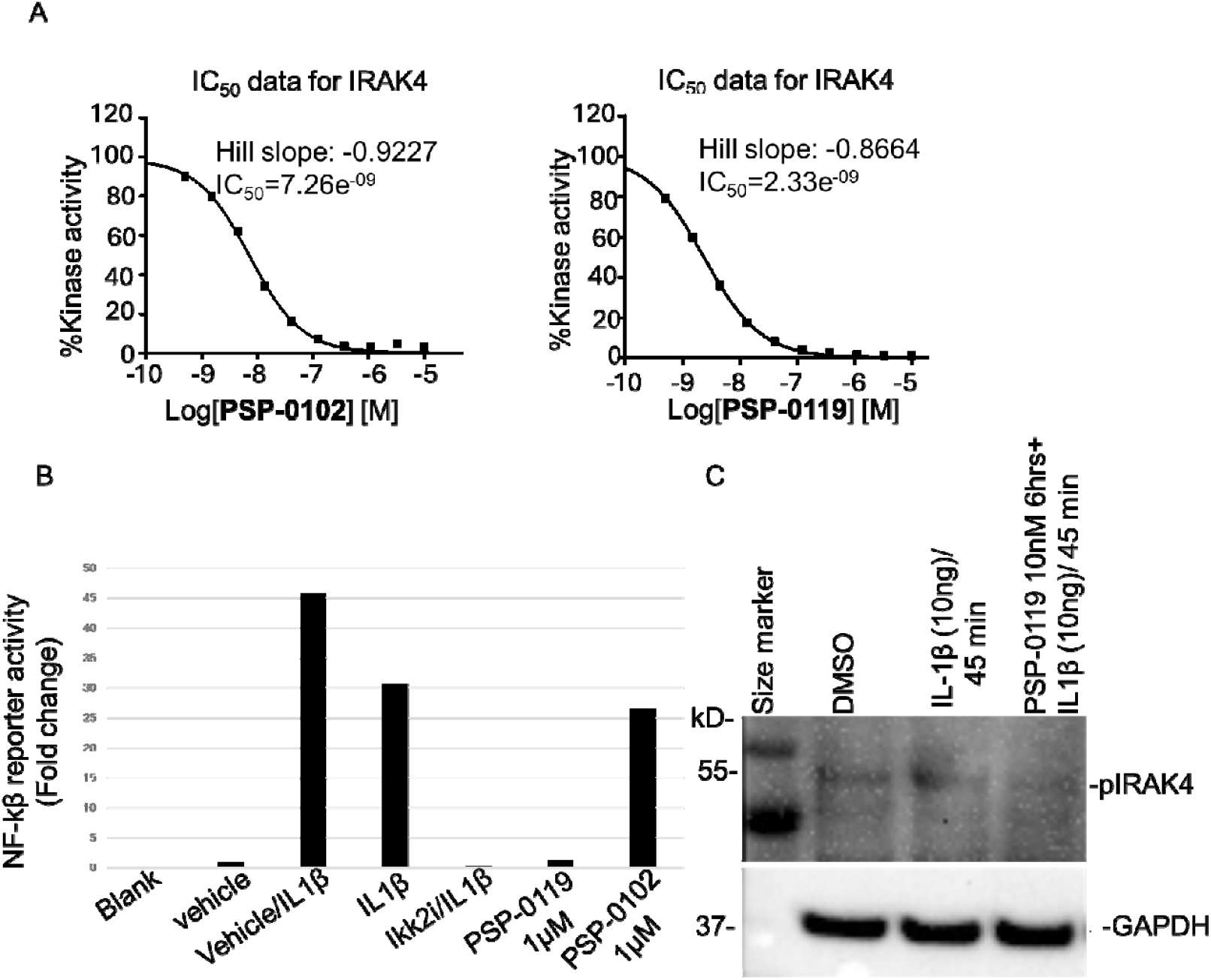
(**A**): Hotspot kinase assay shows that despite drastic modifications on UR241-2, the kinase inhibition activity of PSP-0102 and PSP-0119 was not lost and showed IRAK4 kinase inhibition at 2.33nM. (**B**): PSP-0119 inhibited NF-kβ reporter activity induced by IL1β. (**C**): PSP-0119 pretreatment blocked the IL1β induced IRAK4 phosphorylation.

**Figure-3.**
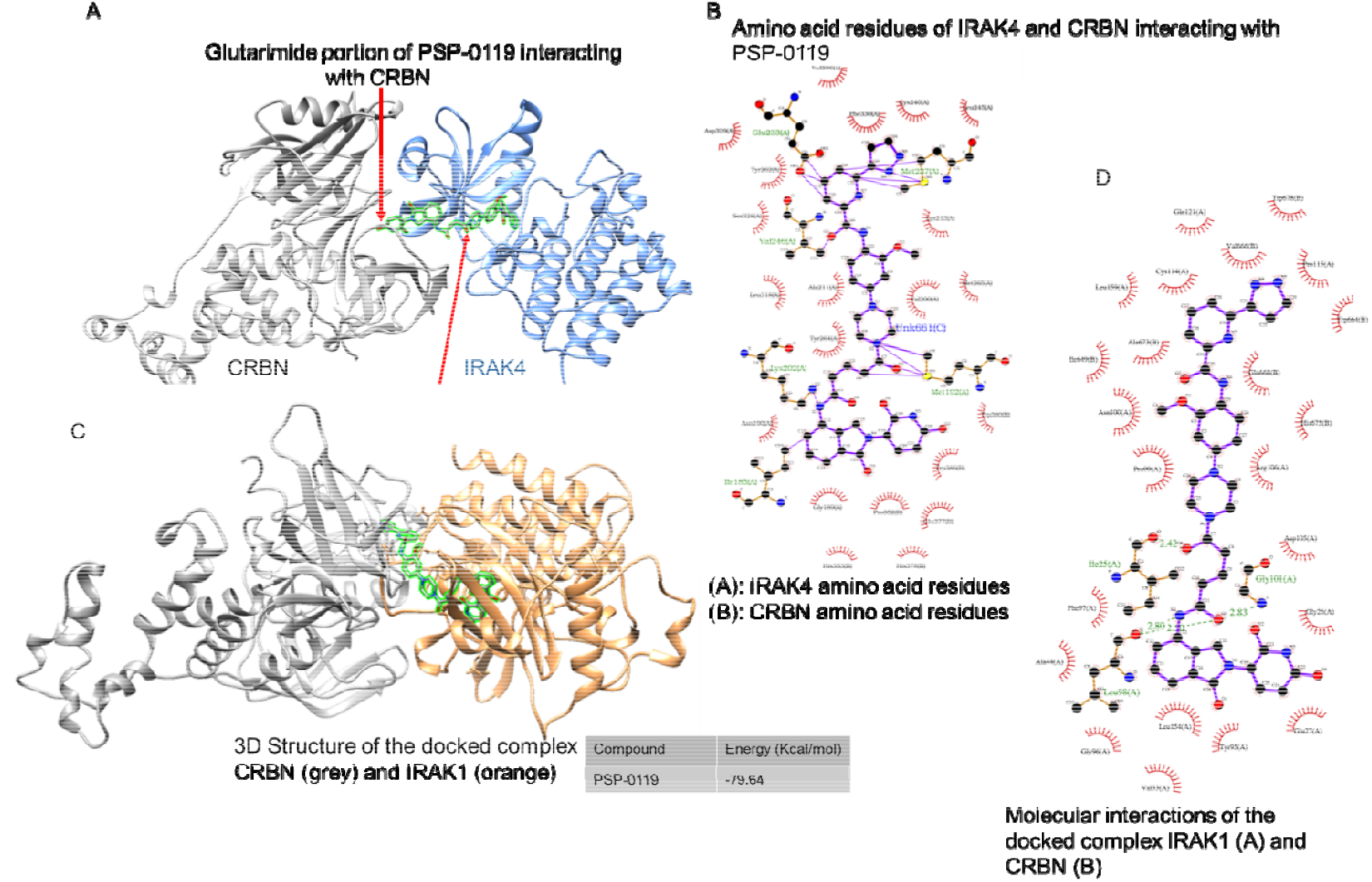
*In silico* docking of PSP-0119 and PSP-0102 with CRBN and IRAK1 and IRAK4 complex. (A): *In silico* docking showed how glutarimide ring of lenalidomide interacts with CRBN and how pyrazolyl-pyridinyl portion engages with IRAK4. Red arrows point to both interactions. (B): Individual CRBN and IRAK4 residues interacting with PSP-0119 atoms are shown. (C): *In silico* docking showed how glutarimide ring of lenalidomide interacts with CRBN and how pyrazolyl-pyridinyl portion engages with IRAK1. (D): Individual CRBN and IRAK1 residues interacting with PSP-0102 atoms are shown.

Next, we analyzed the IRAK4 degradation potential of PSP-0119 utilizing a panel of THP1 (FLT3-wt), MOLM-13, and MV-4-11, FLT-3-ITD-mutant cell-lines. Immunoblotting (raw data: Supplementary Figure-S2) showed that PSP-0119 did not downregulate IRAK4 expression in THP1 cells until the tested doses of 5µM (Figure-4A) whereas between doses 20-40nM doses, IRAK4 expression in FLT3-mutant MV-4-11 and MOLM-13 cells was reduced within 24 hrs of treatment (Figure-4D and E). DC50 calculations showed degradation at 29.12 and 31.56nM for MV-4-11 and MOLM-13 cells respectively (Figure-4F). We confirmed the selectivity of PSP-0119 further using the AML and normal bone marrow (NBM) cells isolated from a FLT-3 wt (Figure-4B) patient and a donor bone marrow (Figure-4C). Immunoblotting of AML patient and NBM cells treated with PSP-0119 showed no degradation of IRAK4 confirming the selectivity towards FLT3 mutant cells (Fig-4B and C). Next, degradation of IRAK1 in MOLM13 and MV-4-11 cells was examined after treatment with PSP-0119. While IRAK1 was robustly degraded in MV-4-11 at ∼10-fold DC_50_ than IRAK4 (Figure-4H), PSP-0119 treatment did not lead to degradation of IRAK1 in MOLM-13 AML cells (Figure-4G). While both IRAK1 and IRAK4mRNA expression both strongly correlate with FLT3mRNA in AML patient’s microarray samples (Figure-5), it is unclear why IRAK1 remains undegraded by PSP-0119 in MOLM-13 cells. One possibility is that PSP-0119 failed to tag IRAK1 in MOLM-13 cells leading to impaired degradation.

**Figure-4.**
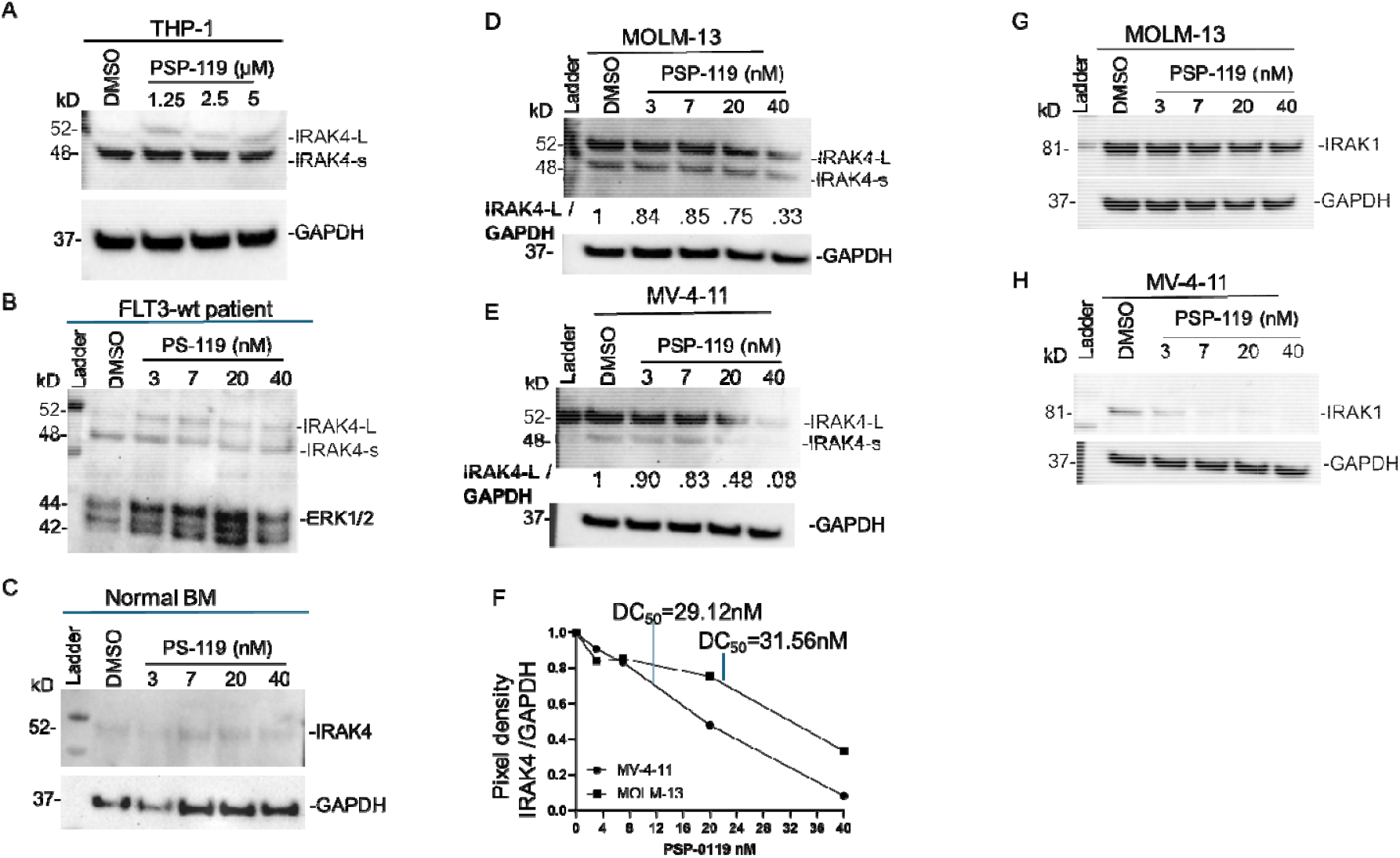
Repeated experiments showed PSP-0119 treatment did not degrade IRAK4 in THP-1 (FLT-3 wild-type) (**A**), FLT-3 wild-type AML patient (**B**) and normal bone marrow cells (**C**). In immunoblotting of FLT-3 normal AML patients (**C)**, total ERK1/2 was used as a loading control as Actin and GAPDH showed variabilities. (**D**): PSP-0119 treatment degraded IRAK4 in MOLM-13 and MV-4-11 (**E**) FLT-3 mutant AML cells between doses 20-40nM during 24 hrs. of treatment (**D** and **E**). Normalized (IRAK4/GAPDH) pixel density is written in numbers in between the bands. (**F**): IRAK4/GAPDH normalized curves and calculated DC_50_ of PSP-0119 against MV-4-11 and MOLM-13 are shown. (**G**): PS-0119 did not degrade IRAK1 in MOLM-13 cells. (**H**): PS-0119 strongly degraded IRAK1 in MV-4-11 cells.

**Figure-5.**
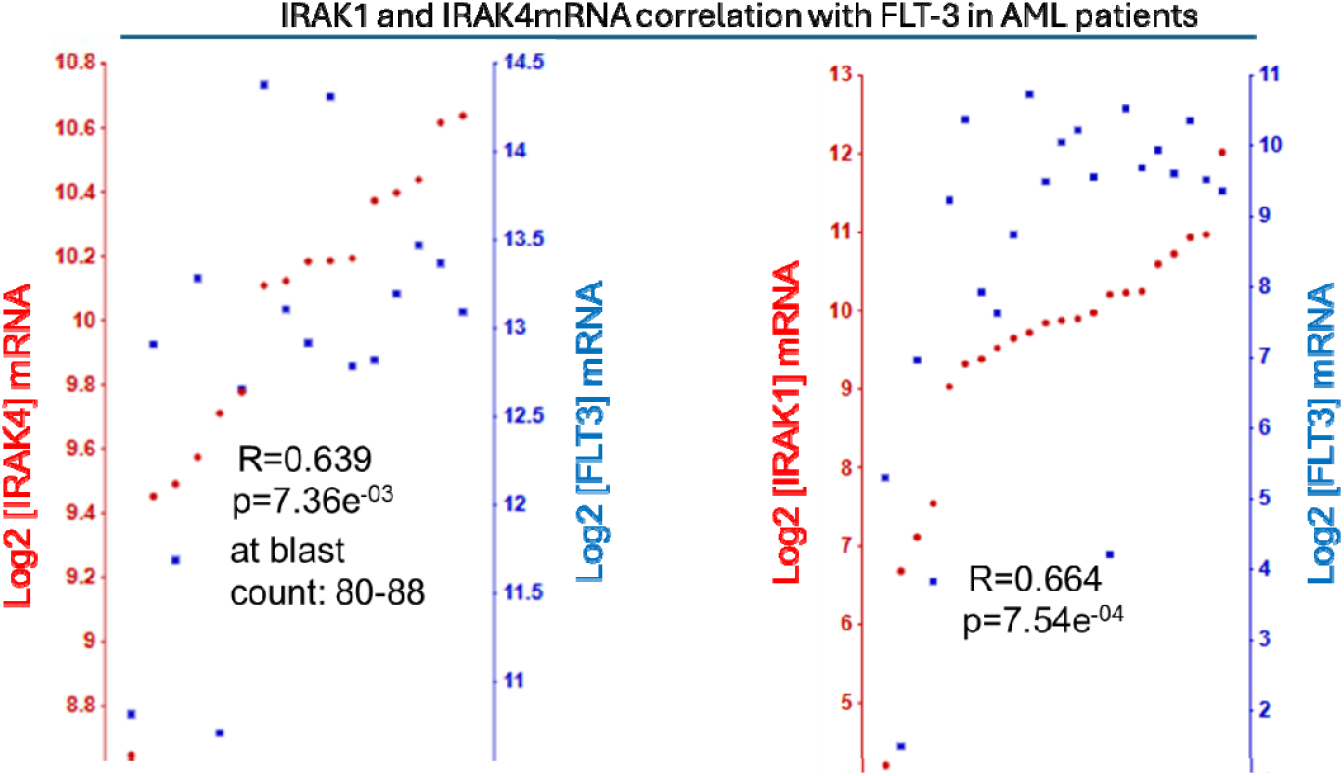
IRAK1mRNA and IRAK4mRNA shows strong correlation with expression of FLT- 3mRNA. Tumor AML 22-MASS.0-u133p2 database (22 samples) were analyzed by the correlation tools available at https://hgserver1.amc.nl/cgi-bin/r2/main.cgi

Next, we examined the effect of PSP-0119 on the global gene expression in MOLM-13 AML cells using bulk-sequencing techniques. In Figure-6A-B (Supplementary Figure-S3), a heatmap of the most altered genes (overexpressed or suppressed) in MOLM-13 after PSP-0119 treatment are shown. Heatmap showed robust downregulation of ENOS-1 (alpha-enolase), an enzyme crucial for cellular metabolism, particularly in glycolysis which is preferred by tumor cells. Analysis of the microarray (Bohlander-422-Mass.0-u133a) using Bloodspot tools showed that ENOS-1 predicted significantly poor prognoses (p=0.0123) (Figure-6C). Next, we examined the effect of PSP-0119 on colony formation (CFU) potential of MOLM-13 and MV-4-11 cells. PSP-0119 treatment reduced CFUs formed by MV-4-11 (Figure-7A); however, repeated attempts with varying cell number seeded and days observed, the CFUs formed by PSP-0119 were statistically insignificant (data not shown). Next, we estimated the viability of THP1, MOLM-13 and MV-4-11 cells upon treatment with PSP-0119 (0, 2.5 and 5μM). PSP-0119 did not affect the viability (measured in terms of mitochondrial metabolism via MTS assay) of FLT-3 wt THP1 cells but FLT-3 mutant MOLM-13 and MV-4-11 cells showed reduced viability during 48 hrs. of treatment (Figure-7B).

**Figure-6.**
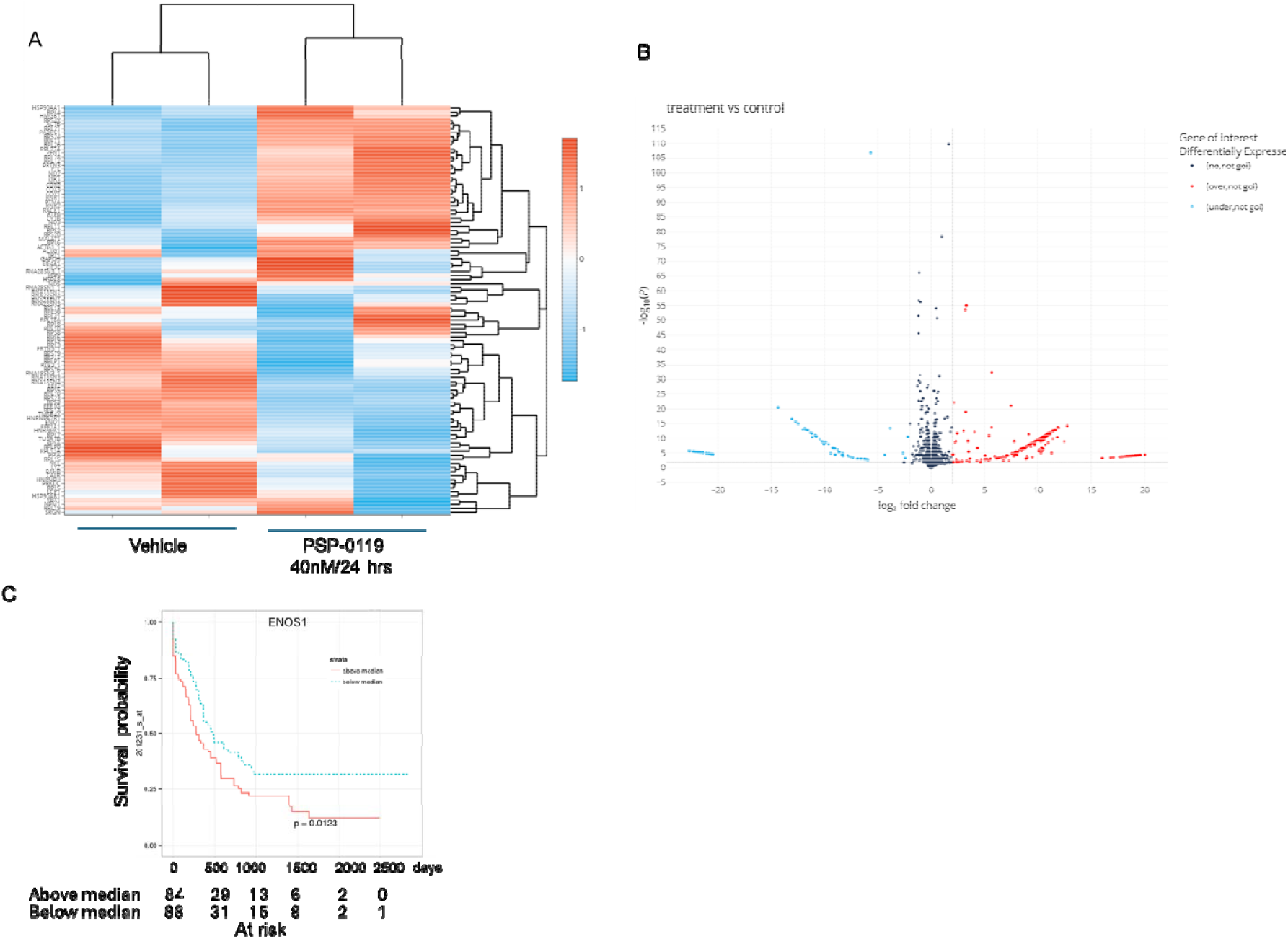
(**A-B**): Bulk-sequencing of PSP-0119 treated MOLM-13 cells have identified gene affected by IRAK4 degradation. (**C**): Volcano plot of the upregulated and downregulated gene after IRAK1/4 degradation in MOLM-13 cells treated with PSP-0119 compared to vehicle. PS-0119 treatment led to downregulation of metabolic driver ENOS-1 gene in MOM-13 cells. Analysis of AML patient’s microarray data available at R2-Genomics and Visualization platform showed that ENOS-1 overexpression predicted poor prognoses in AML patients.

**Figure-7.**
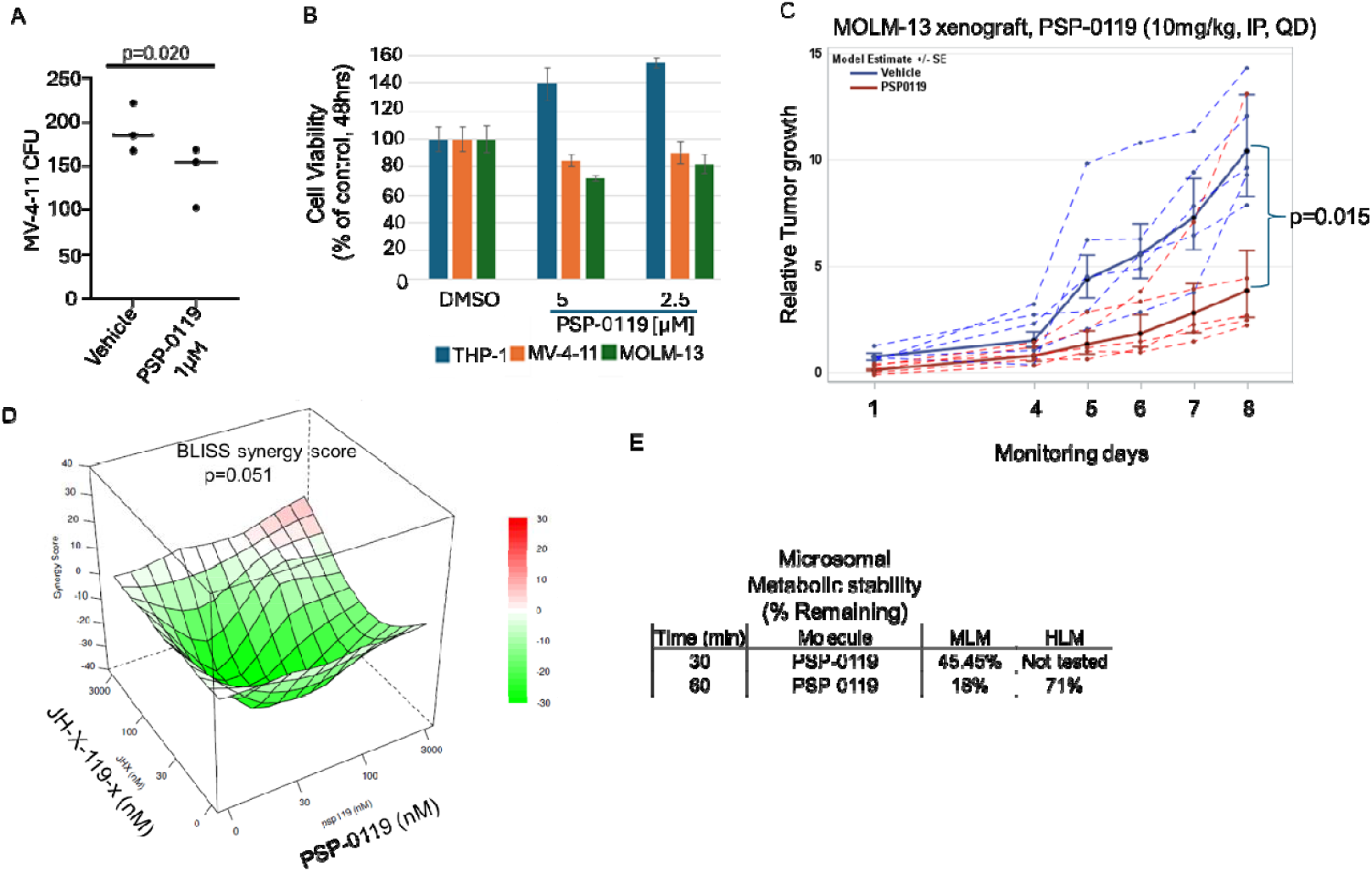
(**A**): PSP-0119 treatment reduced the colonies formed by MV-4-11 AML cells by the 7^th^ day of observation. (**B**): PSP-0119 treatment reduced the viability of MV-4-11 and MOLM- 13 cells. 100,000 cells were seeded in a 96 well plate and treated with indicated concentrations of PSP-0119 for 24 hours. Cell viability was assessed by MTS assay. OD values were read at 490nM. (**C**): PSP-0119 treatment (10mg/kg, IP, once daily/QD, M-F) treatment reduced the growth of MOLM-13 xenograft implanted subcutaneously in NSG mice (p= (1million/mice). All the PSP-0119 treated mice retained normal health and no weight or fur loss, or mobility or socialization disruptions were noted during treatment days. (**D**): The combination of IRAK1 inhibitor JH-X-119-x with PSP-0119 showed synergized reductions of MOLM-13 AML cell- lines viability when measured by MTS assay. (**E**): PSP-0119 showed 18% intact quantity at 60^th^ minute in mice liver microsomes (MLM) but exhibited notable metabolic stability in human liver microsomes (HLM). 71% of PSP-0119 remained intact at 60^th^ minute of observation.

Next, in a pilot experiment, the therapeutic effects of PSP-0119 (20mg/kg, M-F, IP, QD) treatment on the growth of MOLM-13 xenografts implanted subcutaneously in NSG mice were evaluated (raw data, Supplementary Figure-S4).. The initial tumor volume in the treated group was significantly higher than in the vehicle group (Wilcoxon Rank Sum Test p=0.01) (Figure-7C). The estimated tumor volume growth relative to the initial volume for PSP0119 was significantly less than the vehicle group (p=0.015) (Figure-7C). Since combination therapy has consistently shown superior outcomes as compared to monotherapies in AML, we examined the outcome of combining PSP-0119 with IRAK1 inhibitor JH-X-119-01. We observed notable synergy with JH-X-119-01 in MV-4-11 AML cells in vitro (Figure-7D) corroborating literature reports that IRAK1 imparts resistance to IRAK4 inhibition^24^ and blocking both IRAK1 and -4 together impart greater control over AML. Bennett et al have shown that the promising but limited responses to IRAK4 inhibitors in MDS and AML clinical trials are due to functional complementation and compensation by its paralog, IRAK1. IRAK4’s dysregulation has been implicated in various pathological conditions, including cancer^25^, autoimmune disorders, and chronic inflammation, highlighting its potential as a therapeutic target for a diverse variety of diseases^26-2728^. Taken together, degrading IRAK4 via PROTACs emerges as a potential approach to control FLT-3 mutant AML. Next, we examined drug-like characteristics of PSP-0119 using human and murine liver microsomes. HPLC analysis of intact PSP-0119 after 30 and 60 minutes of exposure shows that PSP-0119 carries remarkable stability in human liver microsomes (71%) whereas mouse liver microsomes were able to degrade PSP-0119 by 83%. However, by the 30^th^ minute, 47% of PSP-0119 was found to be intact (Figure-7E).

## Discussions and Conclusion

Ultimately dismal survival outcomes for patients with AML, particularly in the relapsed setting, highlight the urgency to identify novel, druggable driver-like targets, which can be chemically manipulated allowing for development of more effective therapies. We previously examined the evolution of LSC population during patients’ clinical stages and identified a 9-90-fold increase in LSC frequency and expansion of phenotypically diversified LSC population, sustaining AML and pivoting disease recurrence^30^. In these settings, we have previously shown how 1) upregulation of IL1/IL1R Accessory Protein (IL1RAP) signaling in AML^31-33^ and 2) TLR/IL1 agonist signaling, drive bone marrow microenvironment (BMME) remodeling, drive LSC expansion and evolution following treatment and relapse and predict therapy failure in AML^1^, providing the actionable rationale to targeting IL-1/IRAK4 signaling to eradicate LSCs and improve overall AML outcomes. While Anakinra (IL-1-blocker) and Canakinumab (IL-1β-blocker) may carry the potential in treatment of MDS conditions, both have yet to be approved for AML treatment. Previously, we had described a novel small molecule inhibitor of IRAK4 UR241-2 which reduced AML burden in vivo^15^.

In this study, we disclose the outcome of degrading IRAK4 using a novel PROTAC degrader PSP-0119. In addition, we addressed an important question, whether IRAK4, IRAK1, or both should be targeted to control AML. Our CFU assay results in MV-4-11 and MOLM-13 cells indicate that simultaneously targeting both IRAK1 and IRAK4 is more effective in suppressing AML than inhibiting either alone, as these kinases play complementary roles in leukemic cell survival and proliferation through dysregulated innate immune signaling pathways. Treatment with PSP-0119 reduced CFU formation only in MV-4-11 cells, in which IRAK1 was also degraded. In contrast, since PSP-0119 failed to degrade IRAK1 in MOLM-13 cells, it can be inferred that IRAK4 degradation alone was insufficient to suppress CFU formation. IRAK4 inhibition alone may seem less effective due to IRAK1 compensation while, IRAK4 inhibitors (e.g., CA-4948) exhibiting modest responses in AML may partly be sourced compensatory signaling orchestrated by IRAK1 even though it combines the inhibition of FLT3 and CLK. When IRAK4 is inhibited, IRAK1 can maintain leukemic cell function via noncanonical MyD88-independent pathways. FLT-3 is one such non-canonical MYD88-independent factor that IRAK1 and IRAK4 are strongly correlated.

It remains unclarified why FLT-3 mutation is essential for IRAK1 and IRAK4 degradation to occur via PSP-0119. A possibility is mutated FLT3 signaling may enhance the activity of E3 ubiquitin, which tags IRAK1 and -4 with ubiquitin, marking them for degradation in the proteasome. The second possibility is FLT3-ITD hyperactivates downstream pathways including PI3K/ AKT, MAKP and STAT5, which can upregulate specific ubiquitin ligases that target IRAK1 and -4. A third possibility is FLT3 mutation may disrupt normal TLR signaling where IRAK4 is critical for pathway progression. This disruption could lead to feedback mechanisms that downregulate IRAK4 to compensate for hyperactive signaling in leukemic cells. Selective IRAK4 degradation in FLT3 mutant AML cells matters clinically because FLT mutated cells can alter immune signaling and promote AML progression by bypassing or reducing inflammatory checkpoints. This makes IRAK4 a valuable therapeutic target. Stabilizing, or controlling IRAK1/4/FLT3 mutant signaling could restore normal inflammatory or immune responses to slow disease progression. For example, PSP-0119 demonstrated antitumor activity in MOLM-13 FLT3 mutant AML cells derived xenografts backed up with remarkable stability in human liver microsomes. A final question is how to employ PSP-0119 to control AML-type represented by MOLM-13 cell-lines which did not allow IRAK1 degradation (Figure-7A, right). The solution is to combine PSP-0119 with IRAK1 co-valent inhibitor PSP-0119, as the combination of both shows synergistic control over AML cells viability (Figure-7E) validating one of the hypotheses of this study that targeting both IRAK1 and -4 generates superior responses than individual targeting.

## Experimental Methods

### Ethical Use of Animals or Human Participants in Research

Animal uses in this study were approved by IACUC of the University of Rochester (approval no: 2016-011E). Primary AML samples and normal bone marrow (BM) were obtained and characterized as previously described post patient’s consent^15^. We also included nine other AML specimens for immunophenotyping or functional assays. Most of these patients were treated with 7+3 regimen. Bone marrow and peripheral blood (PB) samples from AML patients or normal donors were obtained following informed consent and in accordance with protocols approved by the institutional review board (IRB) at the University of Rochester Medical Center (URMC) and Roswell Park Comprehensive Cancer Center. All samples were processed following the methods outlined in our previous work^34^.

## Experimental procedure

The synthetic route to the target compound PSP-0119 is outlined in Scheme 1. Intermediate 9 was prepared following a previously reported procedure^35^. Similarly, intermediate 8 was synthesized according to an established method with slight modifications^36^. Intermediate 10 was prepared following a previously reported procedure^37^.^3^

### 6-(1H-Pyrazol-5-yl)picolinic Acid (3)

**Step-1:**

A solution of methyl 6-chloropicolinate (2, 3.00□g, 13.04□mmol) in a 4:1 mixture of 1,4-dioxane and water (v/v) was stirred at room temperature under a nitrogen atmosphere. To this, 1- (tetrahydro-2H-pyran-2-yl)-5-(4,4,5,5-tetramethyl-1,3,2-dioxaborolan-2-yl)-1H-pyrazole (1, 5.44□g, 19.56□mmol), K□CO□ (5.58□g, 40.43□mmol), and [Pd(PPh□)□] (0.75□g, 0.65□mmol) were added sequentially. The reaction mixture was heated to 100□°C and stirred for 8□h. Reaction progress was monitored by TLC (ethyl acetate:hexane, 4:6). Upon completion, the mixture was cooled to room temperature, poured into cold water, and extracted with ethyl acetate (3 × 50□mL). The combined organic layers were dried over anhydrous sodium sulfate, filtered, and concentrated under reduced pressure. The crude product was purified by silica gel column chromatography using 15% ethyl acetate in hexane as eluent to afford methyl 6-(1- (tetrahydro-2H-pyran-2-yl)-1H-pyrazol-5-yl)picolinate as a solid (4.51□g, 89% yield).

**Step-2:**

Methyl 6-(1-(tetrahydro-2H-pyran-2-yl)-1H-pyrazol-5-yl)picolinate (4.4 g, 15.33 mmol) was added in a 9:1 mixture of tetrahydrofuran and water (THF:H□O) at room temperature. Lithium hydroxide (LiOH, 0.98 g, 2.33 mmol) was added, and the reaction mixture was stirred at room temperature until complete consumption of the starting material, as monitored by TLC (ethyl acetate:hexane, 1:1, v/v). The reaction mixture was then evaporated to dryness, and the residue was dissolved in water. The aqueous phase was neutralized with citric acid and extracted with ethyl acetate (3 × 50 mL). The combined organic extracts were dried over anhydrous sodium sulfate, filtered, and concentrated under reduced pressure to afford 6-(1-(tetrahydro-2H-pyran-2-yl)-1H-pyrazol-5-yl)picolinic acid as white solid (3.5 g, 83% yield). The crude product was then dissolved in 3% methanolic HCl solution at room temperature by stirring. Stirring continued until the TLC (ethyl acetate:hexane, 7:3, v/v) indicated the complete consumption of starting material. The solvent was evaporated in vacuo and the hydrochloride salt of 6-(1H-Pyrazol-5-yl)picolinic acid (3) was isolated as white solid (2.61 g, 90% yield).

^1^H NMR (400 MHz, CDCl_3_): 8.22 – 8.13 (m, 2H), 8.01 (t, *J* = 7.8 Hz, 1H), 7.73 (d, *J* = 2.2 Hz, 1H), 6.97 (d, *J* = 2.2 Hz, 1H). MS (ESI^+^): calcd for C_16_H_23_N_3_O_5_ = 189.1, found; [M+H]^+^ = 190.1. LCMS purity (%): 99.46 (RT: 0.930).

### Synthesis of tert-butyl 4-(3-methoxy-4-nitrophenyl)piperazine-1-carboxylate (6)

A mixture of compound 4-fluoro-2-methoxy-1-nitrobenzene (4, 1.0 g, 5.8 mmol), tert-butyl piperazine-1-carboxylate (5, 1.09 g, 5.8 mmol), and K□CO□ (0.81 g, 5.8 mmol) was stirred in DMF (10 mL) at 80□°C. The reaction progress was monitored by TLC (ethyl acetate: hexane = 1:1, v/v), which confirmed completion. After cooling to room temperature, the reaction mixture was poured into ice-cold water and extracted with ethyl acetate. The combined organic layers were dried and concentrated under reduced pressure. The crude product was purified by column chromatography (20% ethyl acetate in hexane) to afford intermediate 6 as a yellow solid (1.74 g, 88% yield).

^1^H NMR (400 MHz, DMSO-d_6_): δ = 7.90 (d, *J* = 9.3 Hz, 1H), 6.58 (dd, *J* = 9.4, 2.4 Hz, 1H), 6.53 (d, *J* = 2.4 Hz, 1H), 3.91 (s, 3H), 3.47 (s, 8H), 1.42 (s, 9H). MS (ESI^+^): calcd for C_16_H_23_N_3_O_5_ = 337.1, found; [M+H]^+^ = 338.2. HPLC purity (%): 96.85 (RT: 11.101).

### Synthesis of tert-butyl 4-(4-amino-3-methoxyphenyl)piperazine-1-carboxylate (7)

Compound tert-butyl 4-(3-methoxy-4-nitrophenyl)piperazine-1-carboxylate (6, 1.5 g, 4.4 mmol) was dissolved in methanol (30 mL), and 10% Pd/C was added. The mixture was stirred under a hydrogen atmosphere (20 psi) for 30 minutes. Reaction progress was monitored by TLC (ethyl acetate : hexane = 1:2, v/v), which confirmed completion. The reaction mixture was filtered through a celite pad, and the solvent was removed under reduced pressure. The resulting blue colored product 7 was obtained in 95% yield (1.3 g) and used in the subsequent step without further purification.^1^H NMR (400 MHz, DMSO-d_6_): δ = 6.55 – 6.50 (m, 2H), 6.31 (dd, *J* = 8.4, 2.4 Hz, 1H), 4.26 (s, 2H), 3.74 (s, 3H), 3.46 – 3.40 (m, 4H), 2.90 – 2.85 (m, 4H), 1.41 (s, 9H). MS (ESI^+^): calcd for C_16_H_25_N_3_O_3_ = 307.2, found; [M+H]^+^ = 308.2. HPLC purity (%): 96.27 (RT: 9.356).

### Synthesis of N-(2-methoxy-4-(piperazin-1-yl)phenyl)-6-(1H-pyrazol-5-yl)picolinamide (8)

**Step-1**

To a solution of 6-(1H-Pyrazol-5-yl)picolinic acid (3, 0.3 g, 1.33 mmol) in DMF (5 mL), DIPEA (0.573 mL), EDC·HCl (0.32 g, 1.64 mmol), and HOBt (0.25 g, 1.64 mmol) were added sequentially with stirring. After 15 minutes, tert-butyl 4-(4-amino-3-methoxyphenyl)piperazine-1-carboxylate (7, 0.371 g, 1.20 mmol) was added, and the reaction mixture was stirred at room temperature for 4 hours. Reaction progress was monitored by TLC (ethyl acetate : hexane = 70:30, v/v), which confirmed completion. The mixture was diluted with water (100 mL) and extracted with ethyl acetate. The combined organic layers were dried and concentrated under reduced pressure. The crude product was purified by column chromatography (40% ethyl acetate in hexane) to afford the desired product in 74% yield (0.397 g).

**Step-2**

A solution of compound tert-butyl 4-(4-(6-(1H-pyrazol-5-yl)picolinamido)-3-methoxyphenyl)piperazine-1-carboxylate (0.30 g,0.53 mmol) in 10% TFA in DCM was stirred at room temperature for 3 hours. Reaction progress was monitored by TLC (ethyl acetate : hexane = 70:30, v/v), which indicated complete consumption of the starting material. The solvent was evaporated to dryness, and the residue was washed with hexane. The resulting crude product 13 was obtained in 99% yield (0.200 g) and used directly in the next step without further purification.

^**1**^**H NMR (400 MHz, DMSO-d**_**6**_**) δ** = 10.58 (s, 1H), 9.18 (s, 2H), 8.20 – 8.14 (m, 1H), 8.10 (t, *J* = 7.7 Hz, 2H), 8.06 – 8.01 (m, 1H), 7.86 (d, *J* = 2.0 Hz, 1H), 6.99 (d, *J* = 2.1 Hz, 1H), 6.80 (d, *J* = 2.3 Hz, 1H), 6.62 (dd, *J* = 8.8, 2.3 Hz, 1H), 3.96 (s, 3H), 3.43 – 3.38 (m, 4H), 3.28 – 3.19 (m, 4H). MS (ESI^+^): calcd for C_20_H_22_N_6_O_2_ = 378.2, found; [M+H]^+^ = 379.3. HPLC purity (%): 96.07 (RT: 9.634).

### Synthesis of N-(4-(4-(5-((2-(2,6-dioxopiperidin-3-yl)-1-oxoisoindolin-4-yl)amino) -5-oxopentanoyl)piperazin-1-yl)-2-methoxyphenyl)-6-(1H-pyrazol-3-yl)picolinamide (PSP-0119)

To a solution of compound 5-((2-(2,6-dioxopiperidin-3-yl)-1-oxoisoindolin-4-yl)amino)-5- oxopentanoic acid (9, 0.216 g, 0.58 mmol) in DMF, DIPEA (0.276 mL), EDC·HCl (0.151 g, 0.78 mmol), and HOBt (0.121 g, 0.78 mmol) were added successively with stirring. After 15 minutes, compound N-(2-methoxy-4-(piperazin-1-yl)phenyl)-6-(1H-pyrazol-5-yl)picolinamide (8, 0.200 g, 0.52 mmol) was added, and the reaction mixture was stirred at room temperature for 4 hours. Reaction progress was monitored by TLC (ethyl acetate : hexane = 70:30, v/v), which confirmed completion. The reaction mixture was diluted with water (100 mL) and extracted with ethyl acetate. The organic extracts were combined, dried, and concentrated under reduced pressure. The crude product was purified by column chromatography using 5% methanol in dichloromethane as eluent to afford the desired compound in 64% yield (0.25 g).

^**1**^**H NMR (400 MHz, DMSO-d**_**6**_): δ = 13.22 (s, 1H), 11.03 (s, 1H), 10.30 (s, 1H), 9.82 (s, 1H), 8.26 – 8.28 (m, 1H), 8.11 – 8.02 (m, 2H), 7.94 (d, J = 7.6 Hz, 1H), 7.82 (d, J = 6.8 Hz, 1H), 7.52- 7.47 (m, 2H), 6.97 (s, 1H), 6.77 (s, 1H), 6.58 (d, J = 8.8 Hz, 1H), 5.17 – 5.12 (m, 1H), 4.43 – 4.32 (m, 2H), 3.99 (s, 2H), 3.86 (s, 1H), 3.63 (s, 4H), 3.18-3.13 (m, 5H), 2.87 (s, 1H), 2.73 – 2.58 (m, 1H), 2.50 – 2.43 (m, 4H), 2.37 – 2.31 (m, 1H), 2.04-2.01 (m, 1H), 1.90 – 1.85 (m, 2H).**MS (ESI**^**+**^): calcd for C_38_H_39_N_9_O_7_ = 733.3, found; [M+H]^+^ = 734.4. **HPLC purity (%):** 98.45 (RT: 13.096).

### Synthesis of *N*-(4-(4-(2-(4-(4-(2,6-dioxopiperidin-3-yl)phenyl)piperazin-1-yl)acetyl) piperazin-1-yl)-2-methoxyphenyl)-6-(1H-pyrazol-5-yl)picolinamide (PSP-0102)

To a stirred solution of 2-(4-(4-(2,6-dioxopiperidin-3-yl)phenyl)piperazin-1-yl)acetic acid (10, 0.200 g, 0.60 mmol) in DMF (5 mL) were added DIPEA (0.315 mL, 1.8 mmol), EDC·HCl (0.174 g, 0.90 mmol), and HOBt (0.138 g, 0.90 mmol) successively at room temperature. After stirring for 15 minutes, N-(2-methoxy-4-(piperazin-1-yl)phenyl)-6-(1H-pyrazol-5- yl)picolinamide (8, 0.251 g, 0.66 mmol) was added, and the reaction mixture was stirred at room temperature for 4 hours. The reaction progress was monitored by TLC (ethyl acetate/hexane = 70:30, v/v), confirming completion. The mixture was diluted with water (100 mL) and extracted with ethyl acetate (3 × 50 mL). The combined organic layers were dried over anhydrous sodium sulfate, filtered, and concentrated under reduced pressure. The crude product was purified by column chromatography using 5% methanol in dichloromethane as the eluent to afford the desired compound *N*-(4-(4-(2-(4-(4-(2,6-dioxopiperidin-3-yl)phenyl)piperazin-1- yl)acetyl)piperazin-1-yl)-2-methoxyphenyl)-6-(1H-pyrazol-5-yl)picolinamide (PSP-0102) in 70% yield (0.295 g). ^**1**^**H NMR (400 MHz, DMSO-d**_**6**_): δ = 13.22 (s, 1H), 10.76 (s, 1H), 10.59 (s, 1H), 8.25 – 8.18 (m, 1H), 8.11 – 8.02 (m, 2H), 7.93 (s, 1H), 7.04 (d, J = 8.4 Hz, 2H), 6.97 (s, 1H), 6.89 (d, J = 8.4 Hz, 2H), 6.78 (s, 1H), 6.58 (d, J = 8.6 Hz, 1H), 5.76 (s, 1H), 3.99 (s, 2H), 3.85 (s, 1H), 3.74-3.70 (m, 3H), 3.63 (s, 2H), 3.27 (s, 2H), 3.21 (s, 2H), 3.14 (s, 6H), 2.67 – 2.62 (m, 5H), 2.13 – 2.07 (m, 1H), 2.02 – 1.98 (m, 1H), 1.24 (s, 1H). **MS (ESI**^**+**^): calcd for C_37_H_41_N_9_O_5_ = 691.3, found; [M+H]^+^ = 692.4. **HPLC purity (%):** 95.68 (RT: 9.990).

#### Kinase profiling assay

The kinase profiling assay for PSP-0119 involved testing in 10-dose IC_50_ mode with a 3-fold serial dilution, starting at 20 µM. The control compound, staurosporine, was tested in 10-dose IC_50_ mode with a 4-fold serial dilution, starting at 20 µM. Reactions were carried out with 1 µM ATP, and the % enzyme activity was calculated relative to DMSO controls. Curve fits were performed where the enzyme activities at the highest concentration of compounds were less than 65%. This assay was performed in the laboratories of Reaction Biology Inc.

### In silico docking

The protein structure prediction tool SWISS-MODEL, which utilizes homology modeling, was employed to predict the structures of IRAK1 and IRAK4. Although crystal structures are available for these proteins, SWISS-MODEL^38^ was primarily used to fill in missing residues and atoms. Specifically, IRAK1 and IRAK4 were modeled using PDB^39^ entries 6BFN and 8W3X as templates, respectively. High-resolution X-ray crystal structures of CRBN (PDB ID: 6BOY) were used for subsequent structural analyses. Protein–protein docking was performed using the HDOCK^40^ program, which applies a hierarchical docking algorithm incorporating both template-based modeling and ab initio approaches to generate IRAK1–CRBN and IRAK4–CRBN complexes individually. The resulting optimized complexes served as receptors for molecular docking studies with the compound PS-0119, conducted using the MedusaDock program^41-44^. The docking protocol focused on the binding pocket defined by the co-crystallized ligand in the IRAK1 and IRAK4 structures, ensuring the biological relevance of the predicted binding modes. Each compound was subjected to 100 independent docking iterations to thoroughly sample possible binding conformations and enhance the reliability of the results. The resulting docking poses were subsequently clustered by MedusaDock, with the centroid of each cluster selected as the final candidate conformation.

#### Cell Viability

Cell viability of AML (THP-1, MOLM-13 and MV-4-11) cell-lines upon PSP-0119 treatment were estimated via MTS assay as published previously^45^. Briefly, 100,000 cells of each cell-line were seeded in a flat-bottom 96-well ELISA plate in 90μL of RPMI (cat no: RPMI1640) supplemented with 10% heat inactivated FBS (cat no: and 1% strep antibiotic (cat no: xx). DMSO or DMSO in PSP-0119 (10x) adjusted in RPMI media (10μL) media was added from top to make the volume 100μL in each volume and incubated for 44hrs at which time MTS solution in water (5uL) was added to each well and incubated with the drugs for additional 4 hrs. The optical density (OD) of each well was measured at 48 hours using the BioRad iMark spectrophotometer at 490nM wavelength. Averaged O.D. values of wells without cells (blank wells) were subtracted from each well and plotted. The viability of DMSO wells was normalized 100% and the viability of the drug treated wells was calculated with respect to DMSO well using MS excel or GraphPrism V11.0.

#### NF-κβreporter assay

The reporter activity was measured as previously published^15^. Plated 10/000 THP1 -NF-κβ reporter cell cells/50µl media in dark-background 96 well plate. Placed the plate in the incubator for 15-20min. Added PSP-0119 diluted drug in 100% DMSO in cells in the ratio 50µL diluted drug+ 50µL cells and incubated the plate for 30 min at 37°C in an incubator. Added cytokines (hIL-1β, 10ng/mL and incubated plate for 6 hours at 37°C in an incubator. Added 100µL lysis reagent at room temperature and waited at least 3 min to allow complete cell lysis and measure luminescence within 20 min.

#### Colony Formation assay

To examine the impact of PSP-0119 treatment on colony formation units, a CFU-C assay, using MV-4-11 cells was conducted following the published procedure^15^. MV-4-11 cells were plated in methylcellulose-based medium for CFU-C assays (1 × 10^4^ cells/mL. In the suspension, the indicated doses of PSP-0119 were added and mixed very thoroughly and plated in 6-well plates and allowed to incubate thereafter. Colonies were scored after 7 days of culture. Only cell clusters containing more than 50 cells were counted as a colony.

#### Western Blot analysis

To conduct immunoblotting studies, THP-1, MV-4-11, MOLM-13, a FLT3-wt patient donated AML cells and normal bone marrow (BM) cells were cultured in RPMI medium plus 10% heat inactivated FBS (3-4 × 10^6^ cells/mL) containing indicated doses of PSP-0119, as published previously^45^. Cells were collected, washed with cold 1x PBS, and lysed using the lysis buffer (Cell Signaling Tech, cat no:9803) for immunoblotting assays. The PVDF membrane containing the transferred protein was blocked in 5% fat free milk solution for 30 minutes and probed with rabbit IRAK4 antibody (Cell Signaling Tech, cat. no:4363, dilution: 1:1000), rabbit IRAK1 antibody (Cell Signaling Tech, cat. no:4504, dilution: 1:1000). Membranes were striped with OneMinute stripping buffer (GM Biosciences, cat no:GM6001) and were re-probed with rabbit GAPDH antibody (Cell Signaling Tech, cat no: 2118, dilution: 1:10,000). These membranes are preserved.

#### Xenograft assay and Statistical Analyses

Viable MOLM-13 cells (3 million cells/mice) were suspended in cold matrigel:RPMI media (without serum and antibiotics) (1:1) and injected in the right flank of female NSG mice (100μL/mice). When the tumors reached noticeable sizes, mice were randomized in a group of n=5 for control and treatment groups. Mice were treated with vehicle (β-cyclodextrin+Solutol) or vehicle containing PSP-0119 (equivalent to 10mg/kg, IP, daily). Tumor sizes were measured on the days indicated using the digital calipers. We assessed the tumor growth in each animal relative to their own initial volume (Current Volume – Initial Volume) / Initial Volume). A repeated measures analysis of variance was performed on the log2 scale using maximum likelihood estimation with treatment, day, and the interaction between treatment and day as fixed effects with SAS software (v9.4, Cary NC). The correlation of repeated measures on the same animal over time was handled using a Toeplitz covariance within treatment group. Model assumptions were verified graphically. No correlation was found between treatment and day, so the model was simplified. The model estimates of the average relative reduction due to PSP-0119 are 0.31 (95%CI=0.13, 0.70) after back-transformation.

#### Synergy assay

Cell viability of AML (MOLM-13) cell-lines upon PSP-0119 and IRAK1 co- valent inhibitor JH-X-119-x treatment was estimated via MTS assay. Briefly, 100,000 cells of each cell-line was seeded in a flat bottom 96 well ELISA plate in 50μL of RPMI (cat no: RPMI1640) supplemented with 10% heat inactivated FBS (Aventor Seradigm, product no:89510-186, lot no:290M23) and 1% strep antibiotic (Gibco, cat no: 15070063). DMSO or PSP-0119 or JH-I-15 in DMSO (10x) adjusted in RPMI media (50μL) media was added in quadruplicate wells from top to make the final volume 100μL in each well and incubated for 44hrs at which time MTS solution in water (5uL) was added to each well and incubated with the drugs for additional 4 hrs. The optical density (OD) of each well was measured at 48^th^ hours using BioRad iMark spectrophotometer at 490nM wavelength. Averaged O.D. values of wells without cells (blank wells) were subtracted from each well and plotted. The viability of DMSO wells was normalized 100% and the viability of the drug treated wells was calculated with respect to DMSO well using MS excel or GraphPrism V11.0.

#### Stability in liver microsomes

Metabolic stability was assessed using mouse liver microsomes under anaerobic conditions. Test compounds (1 µM) were incubated in 0.1 M potassium phosphate buffer (pH 7.4) containing NADPH (1 mM) as the primary cofactor; UDPGA and PAPS were included where required. Verapamil was used as the reference compound. Anaerobic conditions were maintained in a sealed, nitrogen-flushed chamber with vacuum capability and validated prior to study initiation. Reactions were pre-incubated at 37 °C, 500 rpm for 10 min, initiated with NADPH (12.5 mM stock), and sampled at 30 min. Reaction were terminated by adding three volumes of acetonitrile containing an internal standard, vortexed, and centrifuged (4000 × rcf, 20 min, 4 °C). Supernatants were analyzed by LC–MS/MS to determine the percentage of parent compound remaining.

#### Microarray database analyses

The AML patients microarray databases deposited at R2genomics analysis and visualization platform (https://hgserver1.amc.nl/cgi-bin/r2/main.cgi?open_page=login), GEPIA (http://gepia.cancer-pku.cn/detail.php?gene=IRAK4&clicktag=survival), and GENT(http://gent2.appex.kr/gent2/) were analyzed using the system inbuilt tools. Cutoff of the expression selected by the databases was accepted. Kaplan-Meier analysis was analyzed using these tools with median cut-off selected by the databases as high expressor and low expressor.

## ASSOCIATED CONTENT

### Supporting Information

1. NMR, Mass and HPLC characterization charts of all the compound and intermediates
2. Uncut images of all immunoblots
3. Complete bulk-seq heatmap and volcano plots
4. Raw excel sheet: data of the xenograft tumor sizes measured.

## Author Contributions

NK, CS, JPM, KK: cell culture, cell viability assay, CFU assay and immunoblotting, CR and NAS: microarray data analysis, NAS and SE: independent molecular docking. LE: xenograft. JL: AML patient samples. NVD: supervision of molecular docking. MS: statistical analysis. MK: supervised the synthesis of PSP-0119 and PSP-0102 and helped write the experimental section of the manuscript. BME, RT, KK, NCD, RGM, MWB and JL edited the manuscript. MWB and RGM: partial resources. RKS: conceived the project and designed the PROTACs PSP-0119 and PSP-0102, wrote the manuscript and brought partial resources. The manuscript was written through the contributions of all authors. All authors have given approval to the final version of the manuscript.

## ABBREVIATIONS

AML: Acute myeloid leukemia
CFU: Colony formation units
CRBN: Cereblon
DMSO: Dimethyl sulfoxide
MTS: Methyl triazolium salt
FLT3-ITD: FMS-like tyrosine kinase 3 internal tandem duplication
IRAK1: Interleukin receptor associated kinase-1
IRAK4: Interleukin receptor associated kinase-4
IL-1β: Interlekin-1 beta
MAPK: Mitogen-activated protein kinase
MyD88: Myeloid differentiation primary response 88
MDS: Myelodysplastic syndrome
MPN: myeloproliferative neoplasms
NSG mice: NOD.Cg-Prkdc scid Il2rg tm1Wjl/SzJ mice
NF-κB: IκB kinase (IKK)-nuclear factor-κB,
OD: Optical density
PI-3K: Phosphoinositide 3-kinase
PROTAC: Proteolysis Targeting Chimeras
TLC: Thin layer chromatography

