## Supplemental data for "PSP-0119: Targeted IRAK4 Degradation as a Novel Therapeutic Strategy for FLT3-Mutant AML"

**Supplementary Information**

**Figure S1. ^1^H NMR spectrum of intermediate 3.**

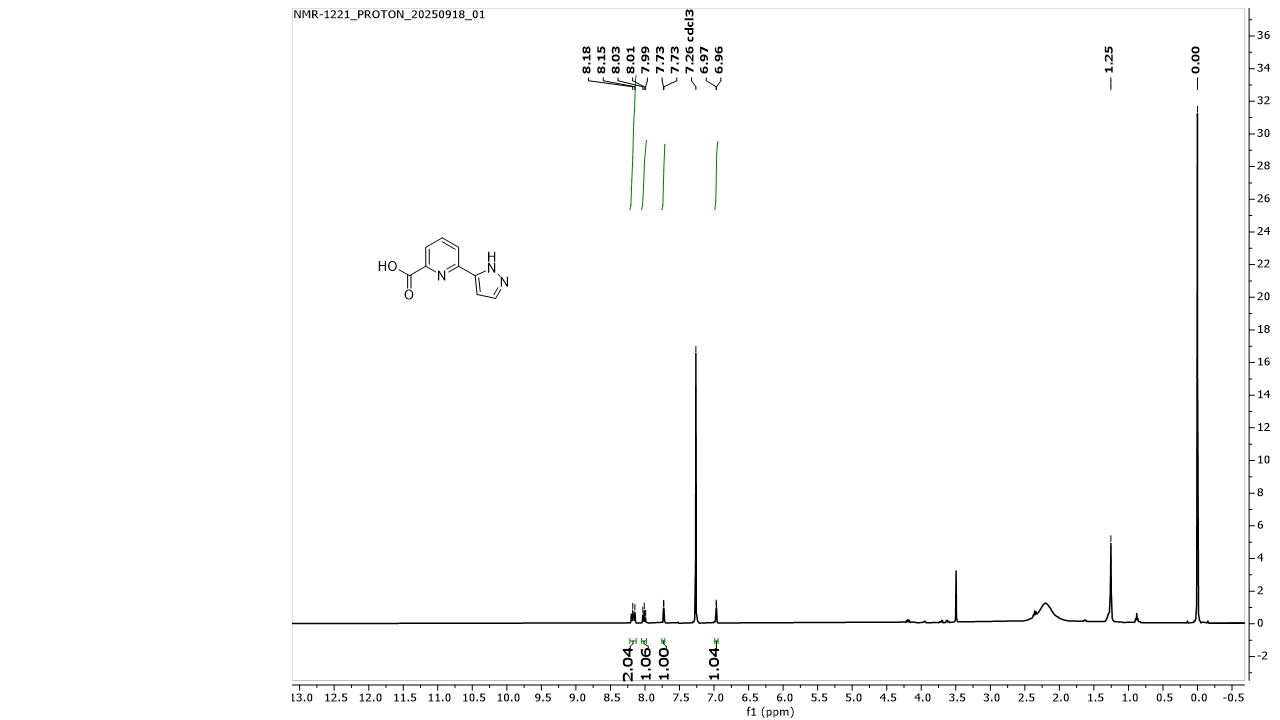

**LC_MS of intermediate 3**

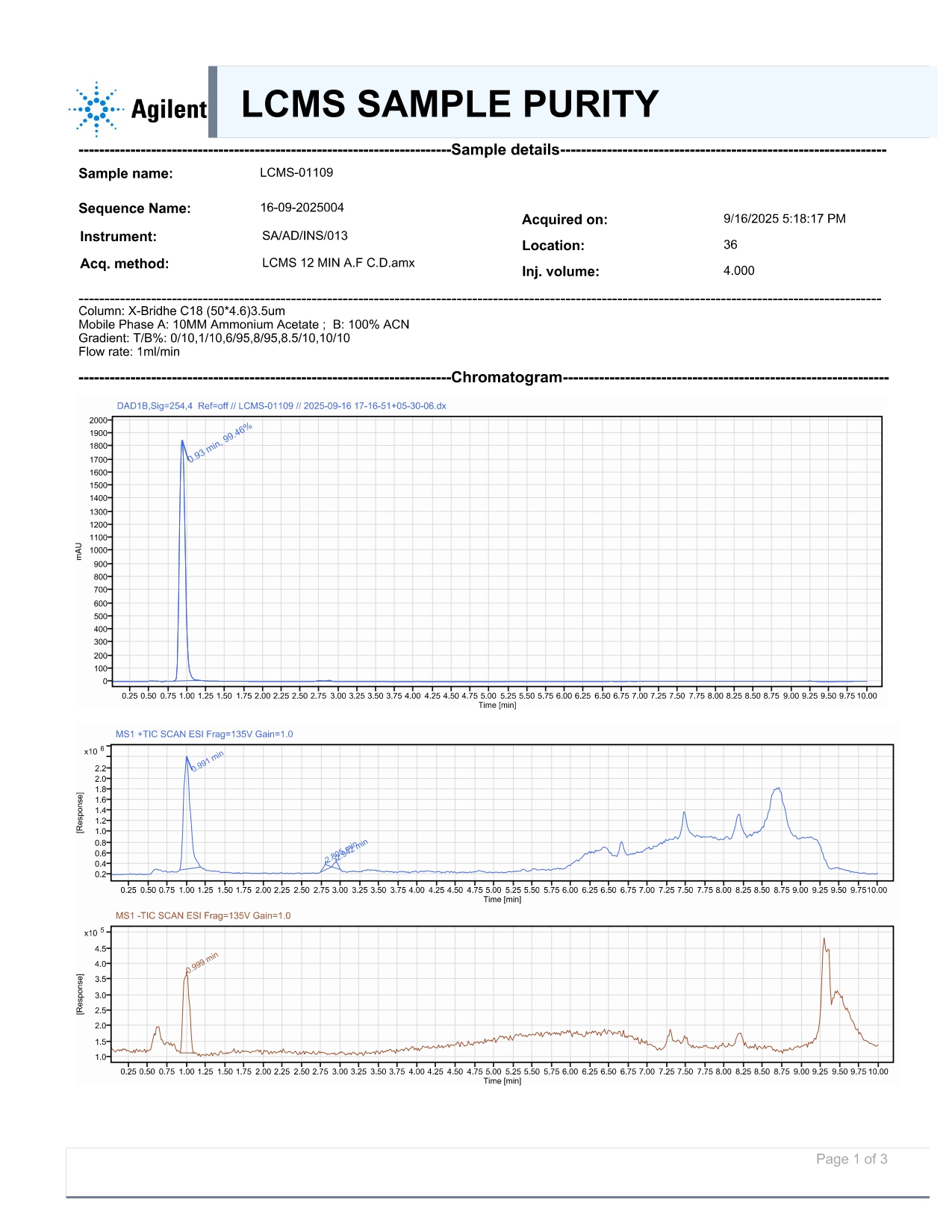

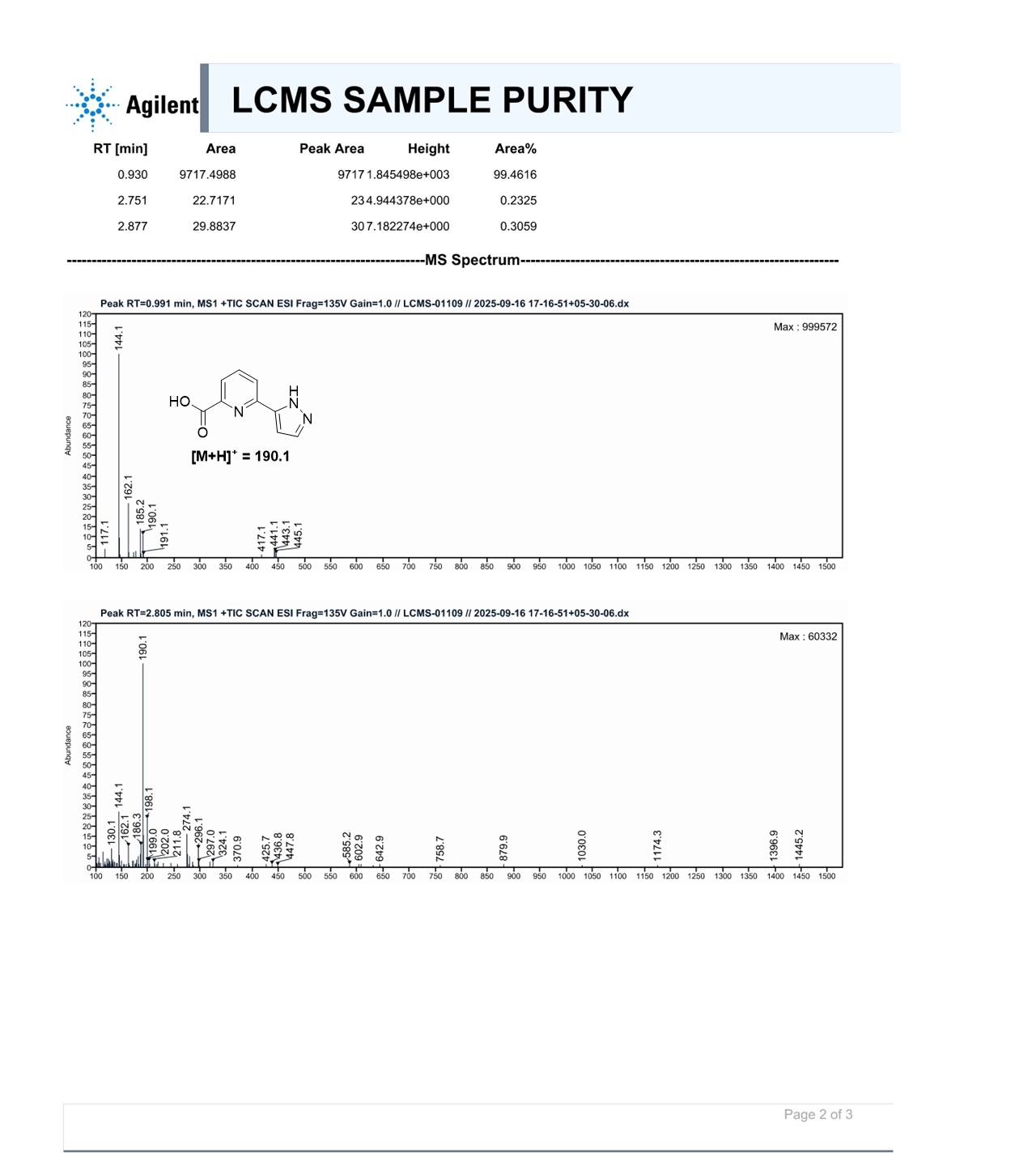

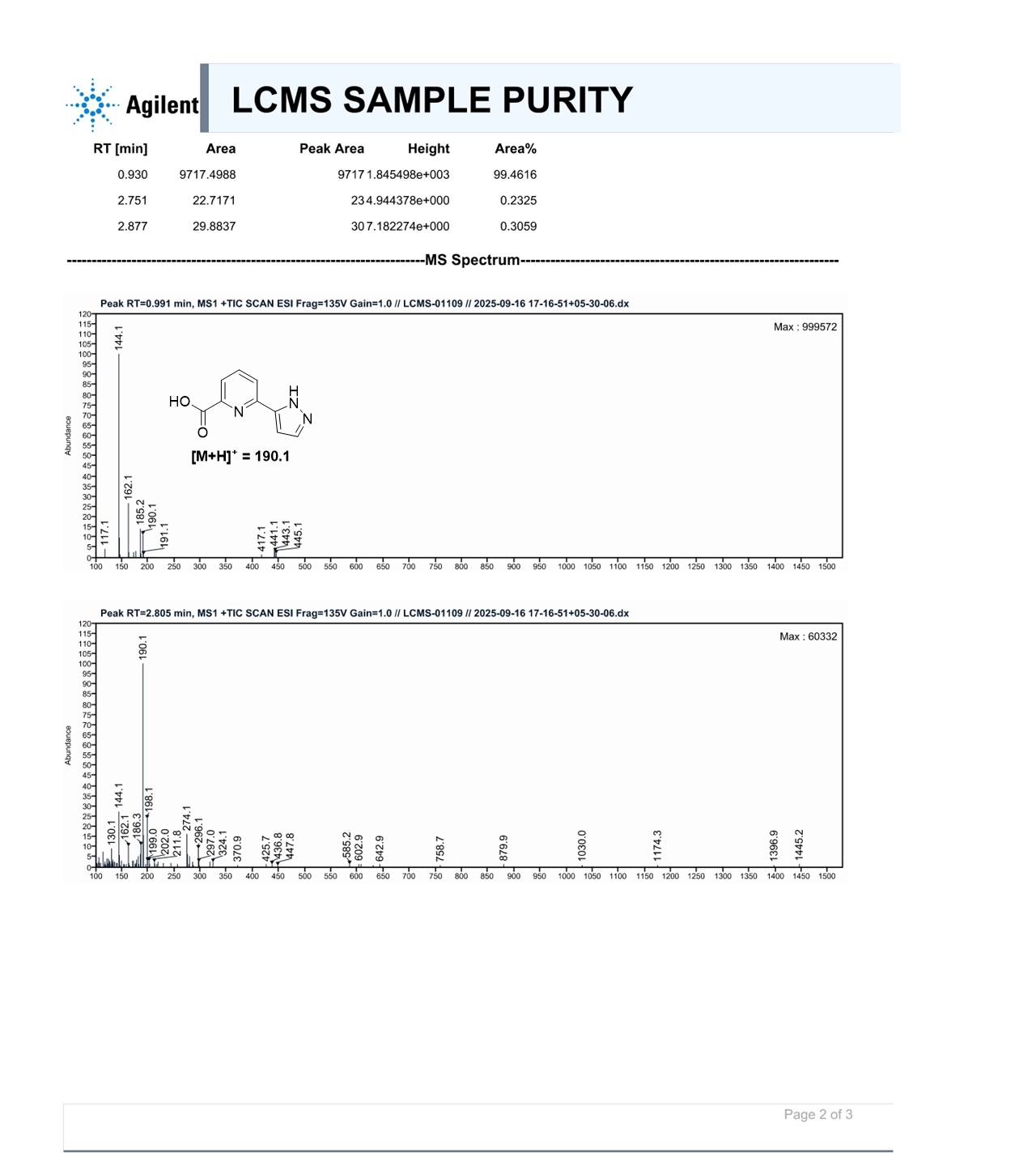

**Figure 2. ^1^H NMR spectrum of intermediate 6.**

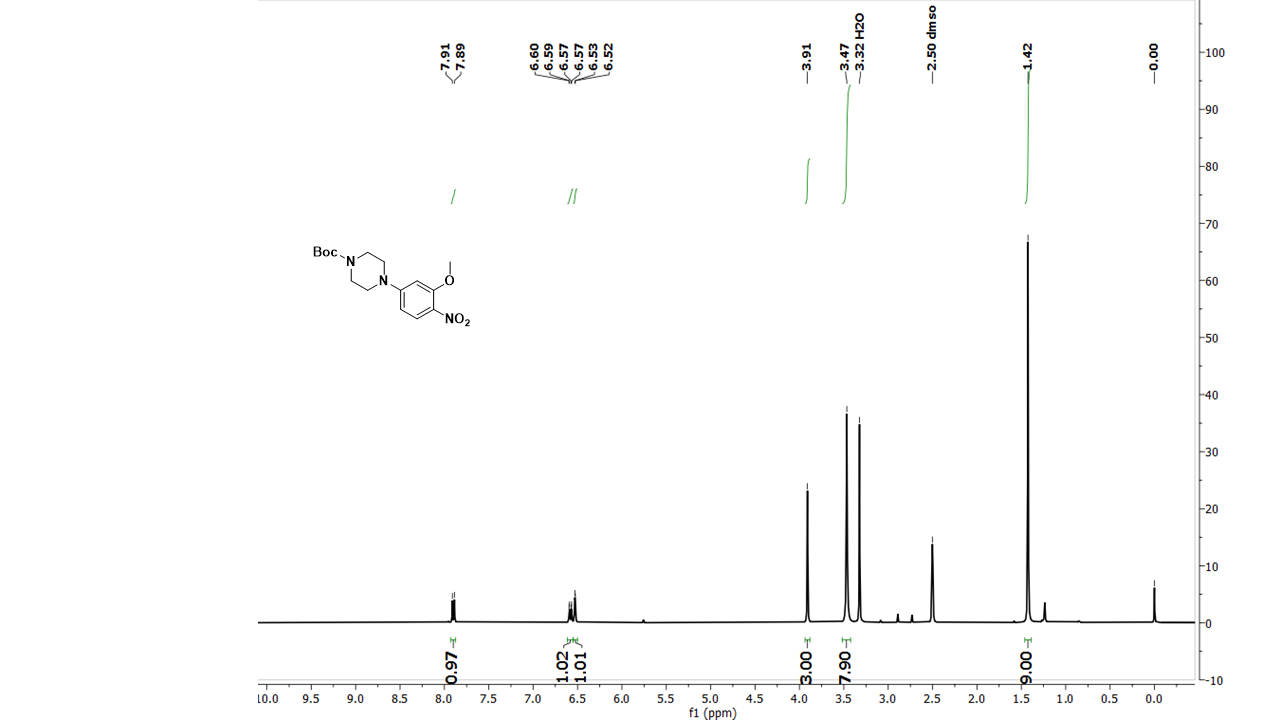

**Figure 3. MS spectrum of intermediate 6.**

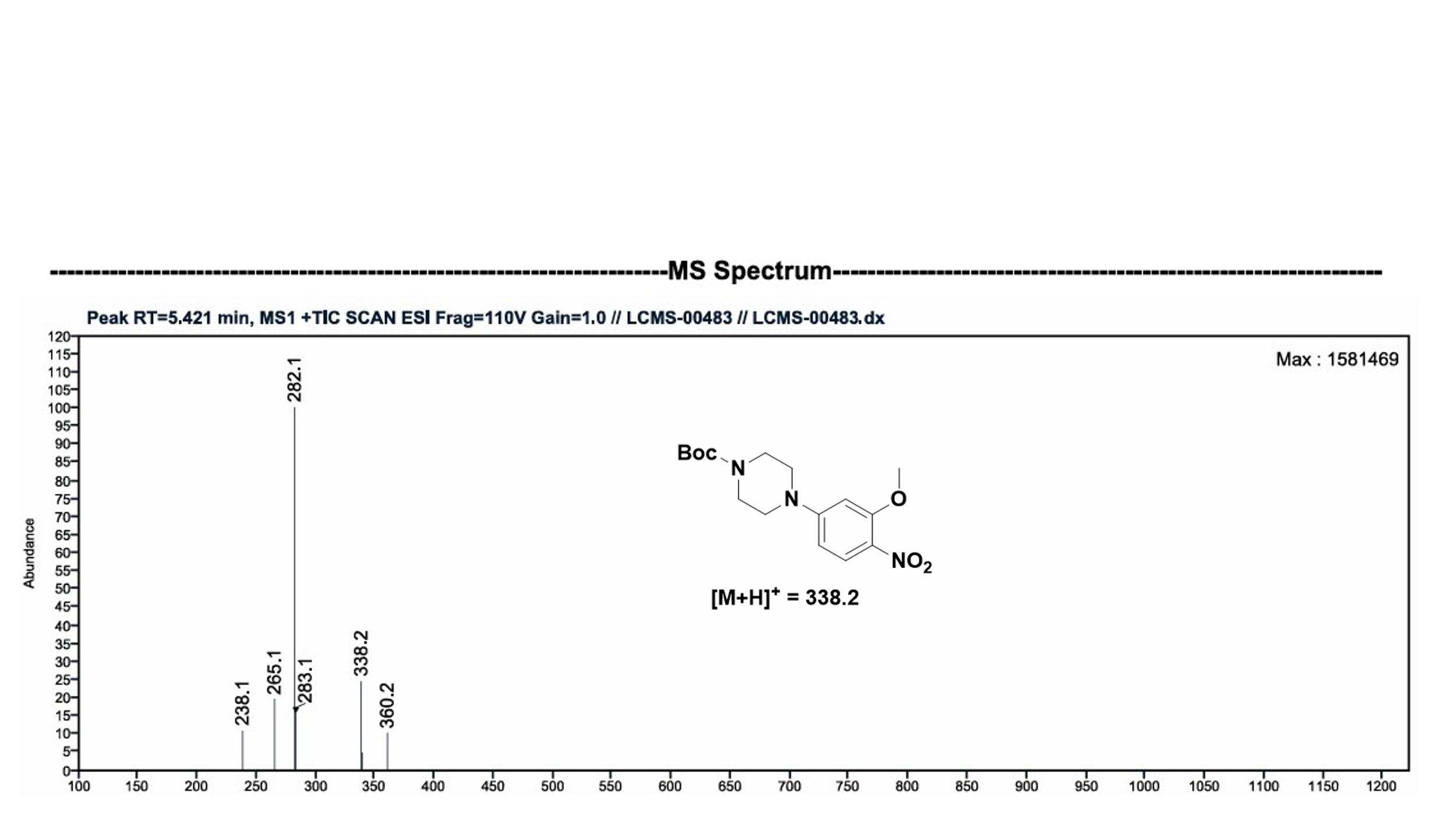

**HPLC of intermediate 6.**

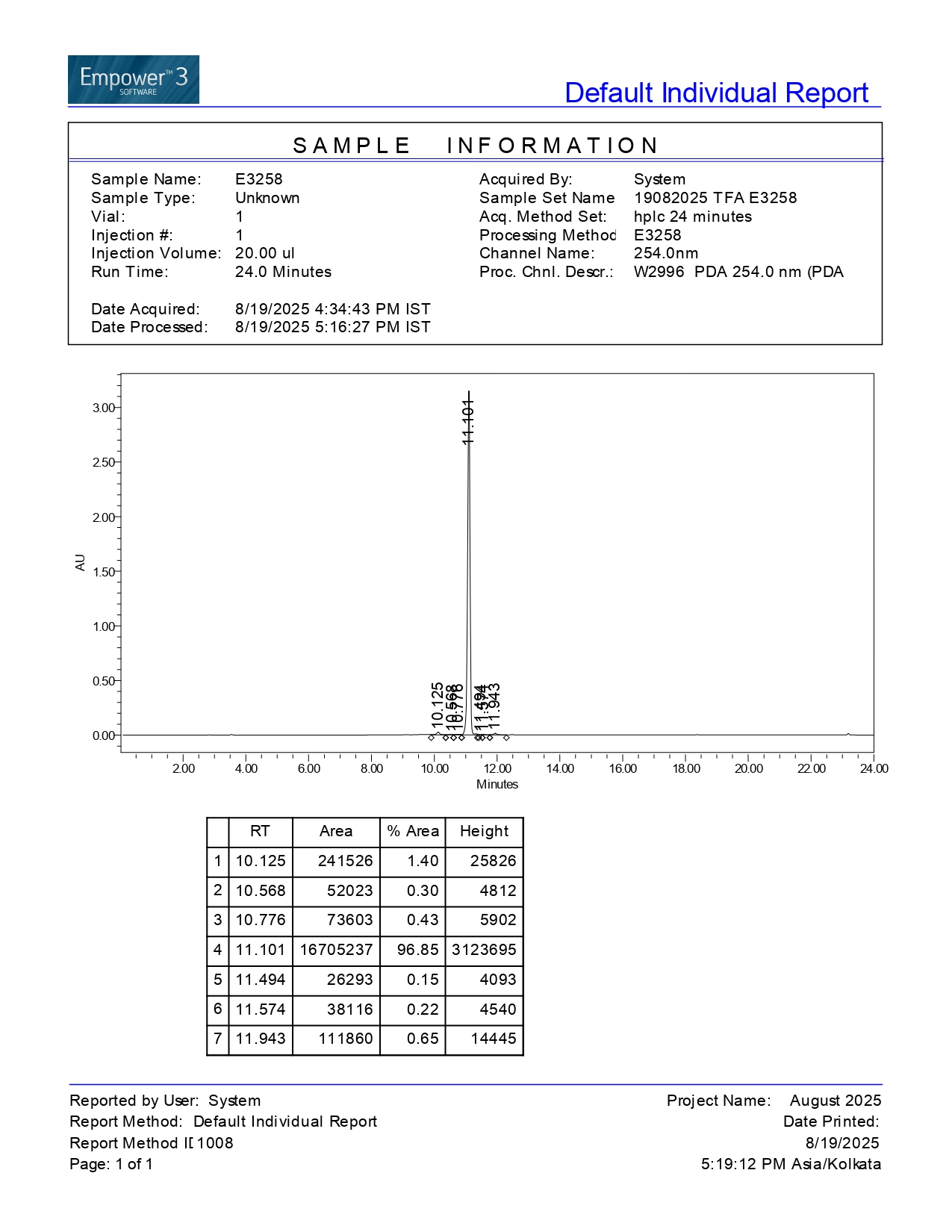

**Figure 4. ^1^H NMR spectrum of intermediate 7.**

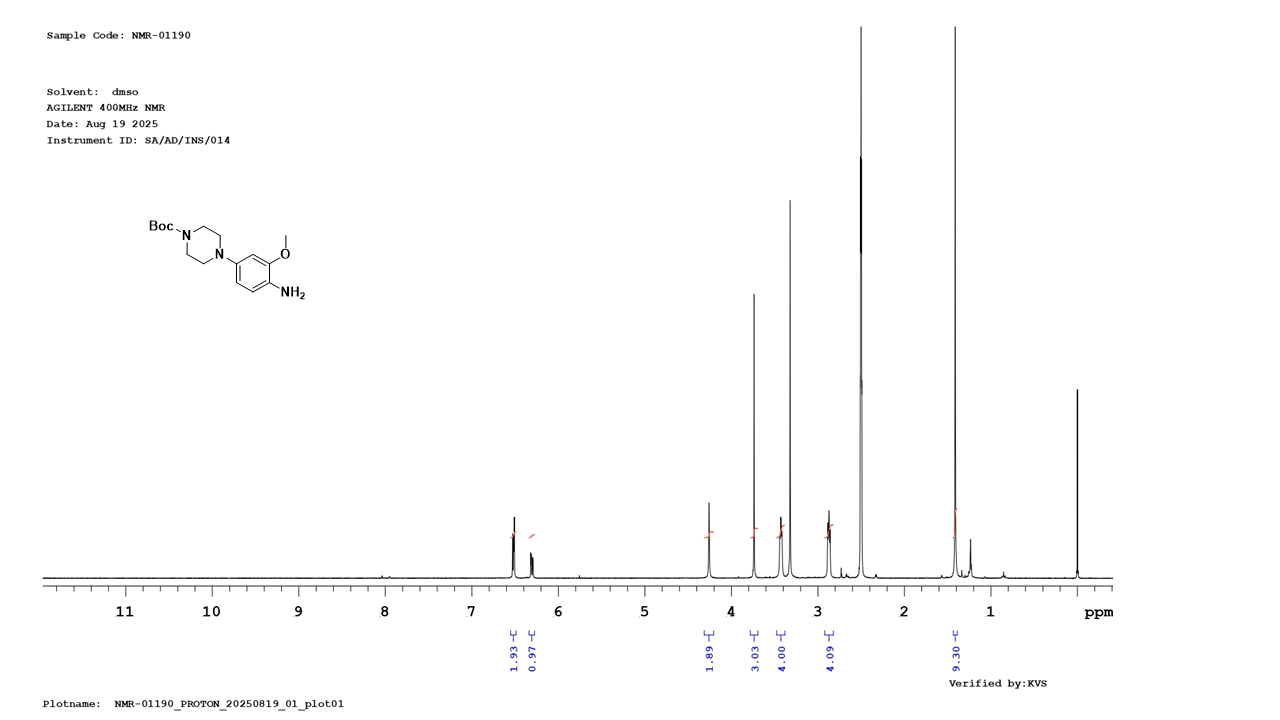

**Figure 5: MS of intermediate 7**

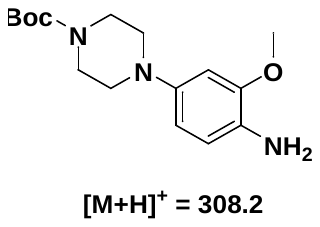

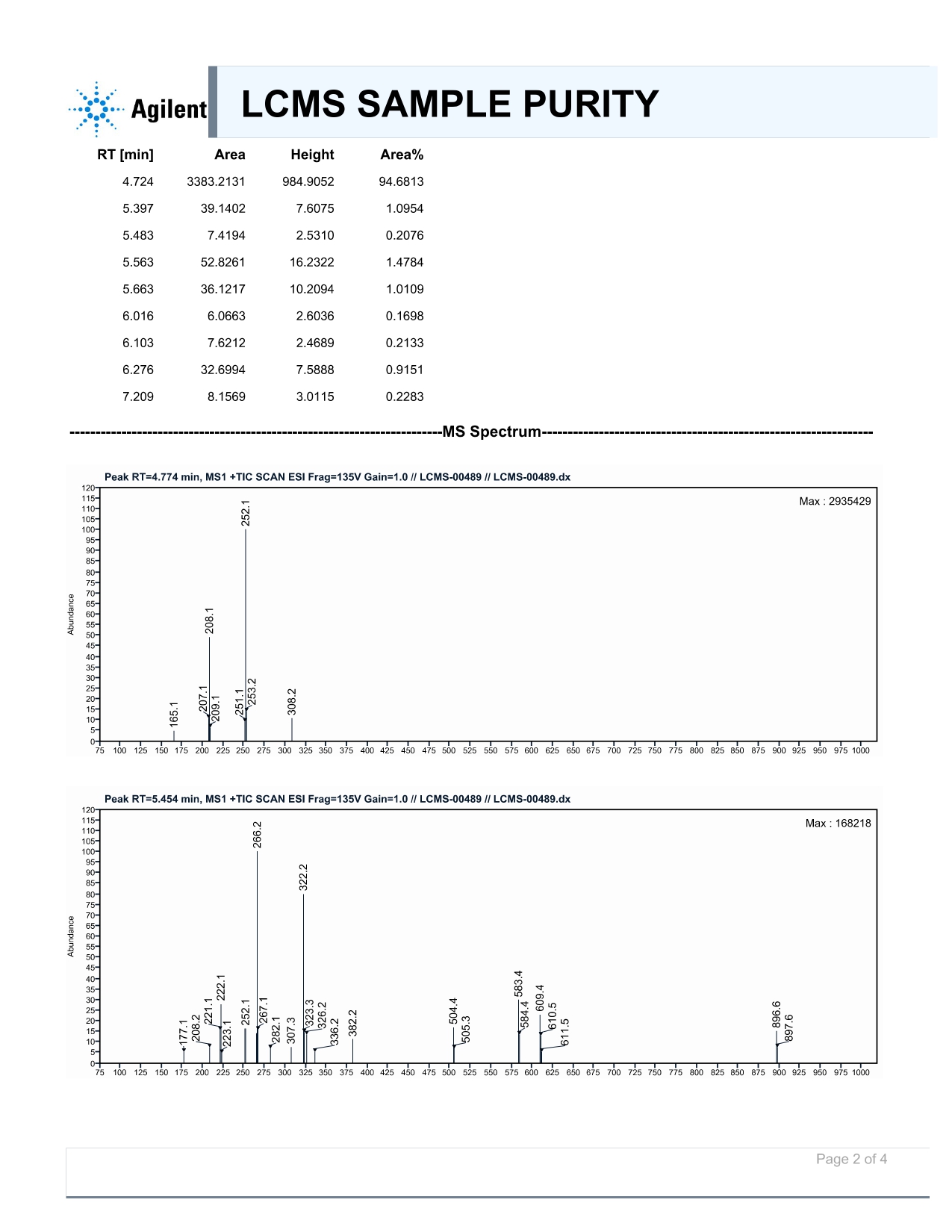

**HPLC of intermediate 7.**

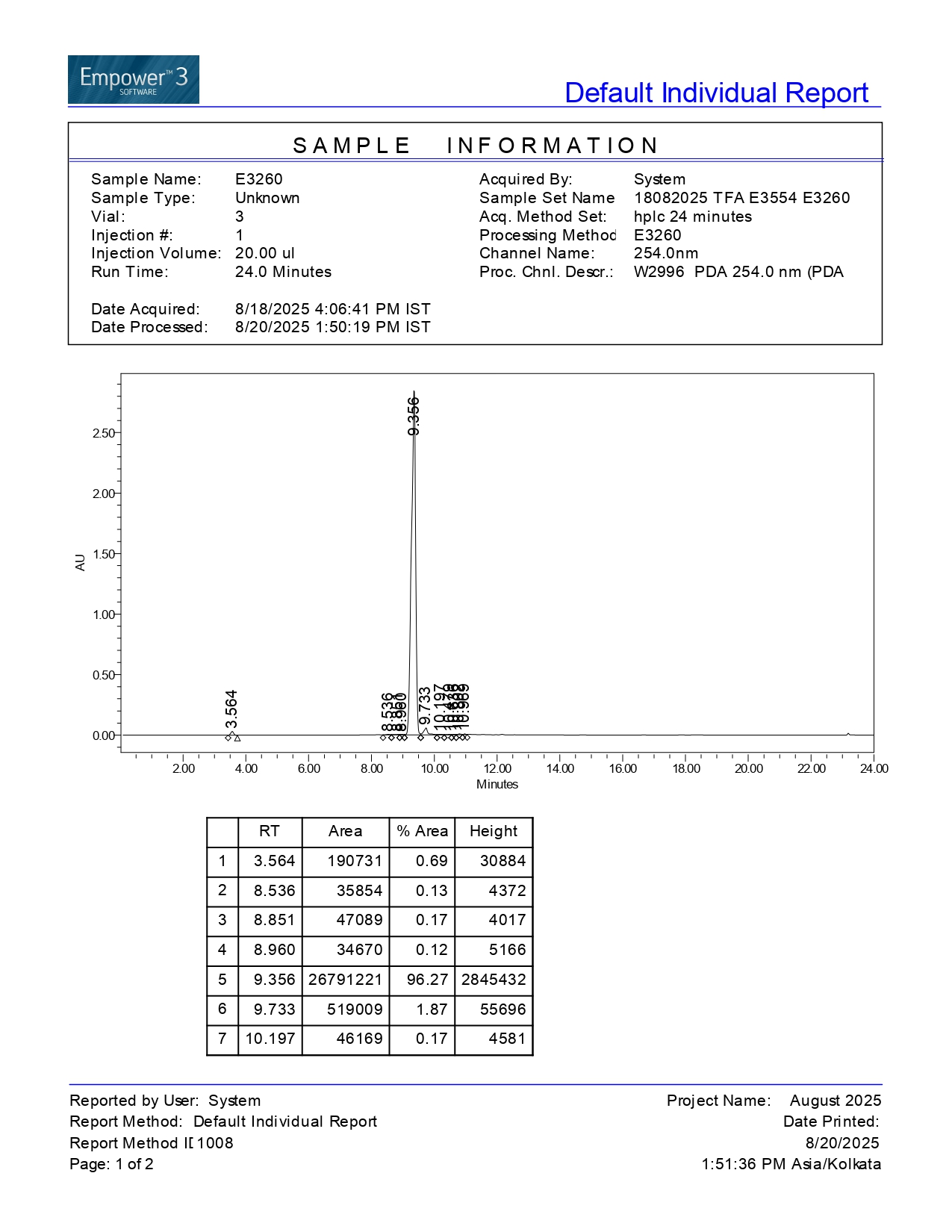

**Figure 6. ^1^H NMR spectrum of intermediate 8.**

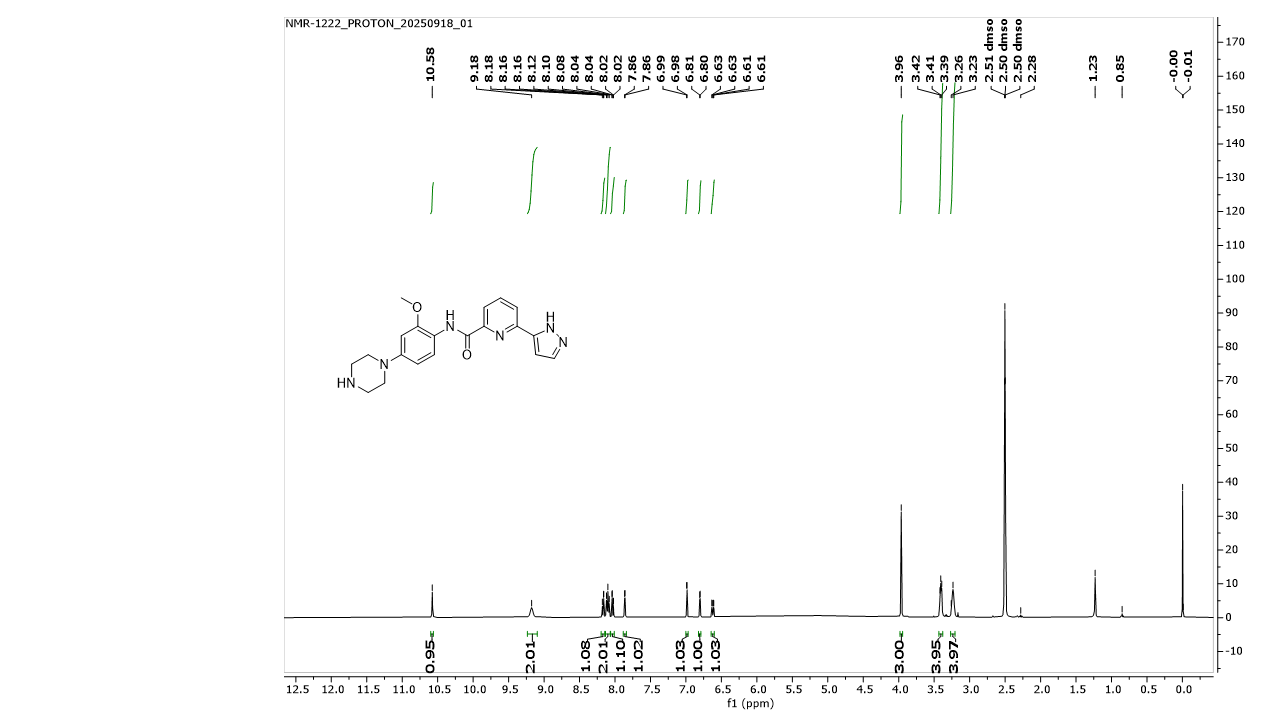

**Figure 7. MS spectrum of intermediate 8.**

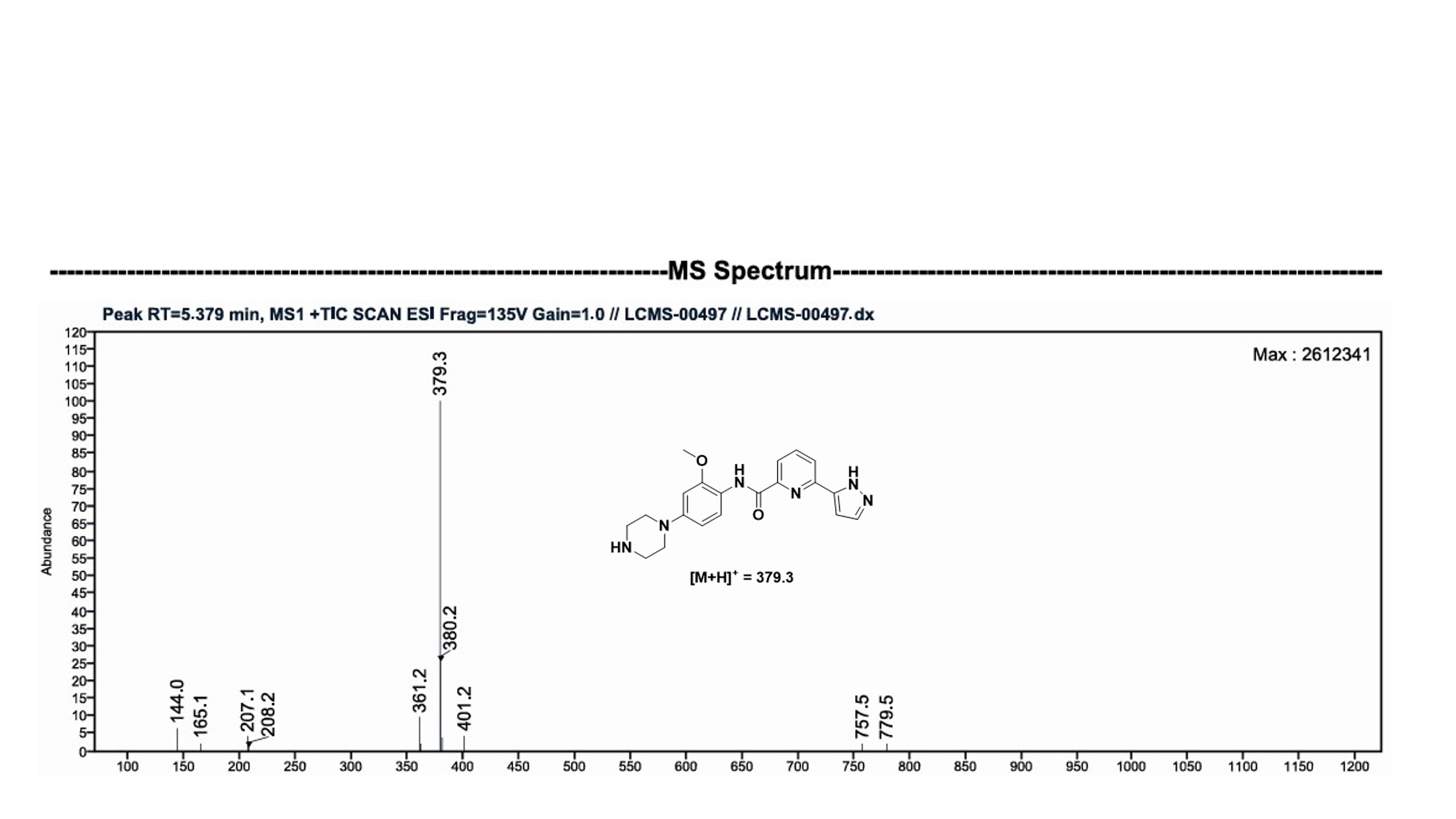

**HPLC of intermediate 8.**

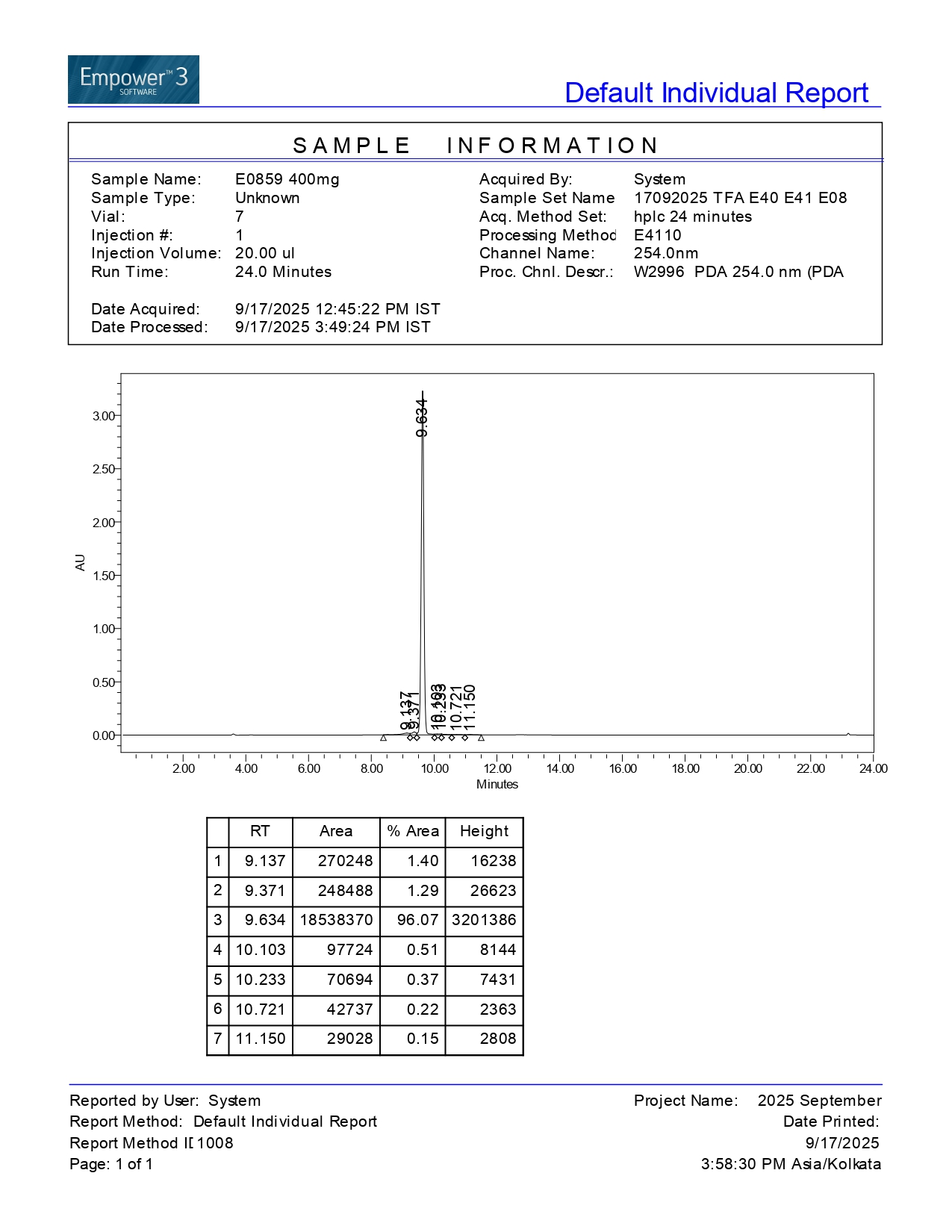

**Figure 8. ^1^H NMR spectrum of intermediate PSP-0119.**

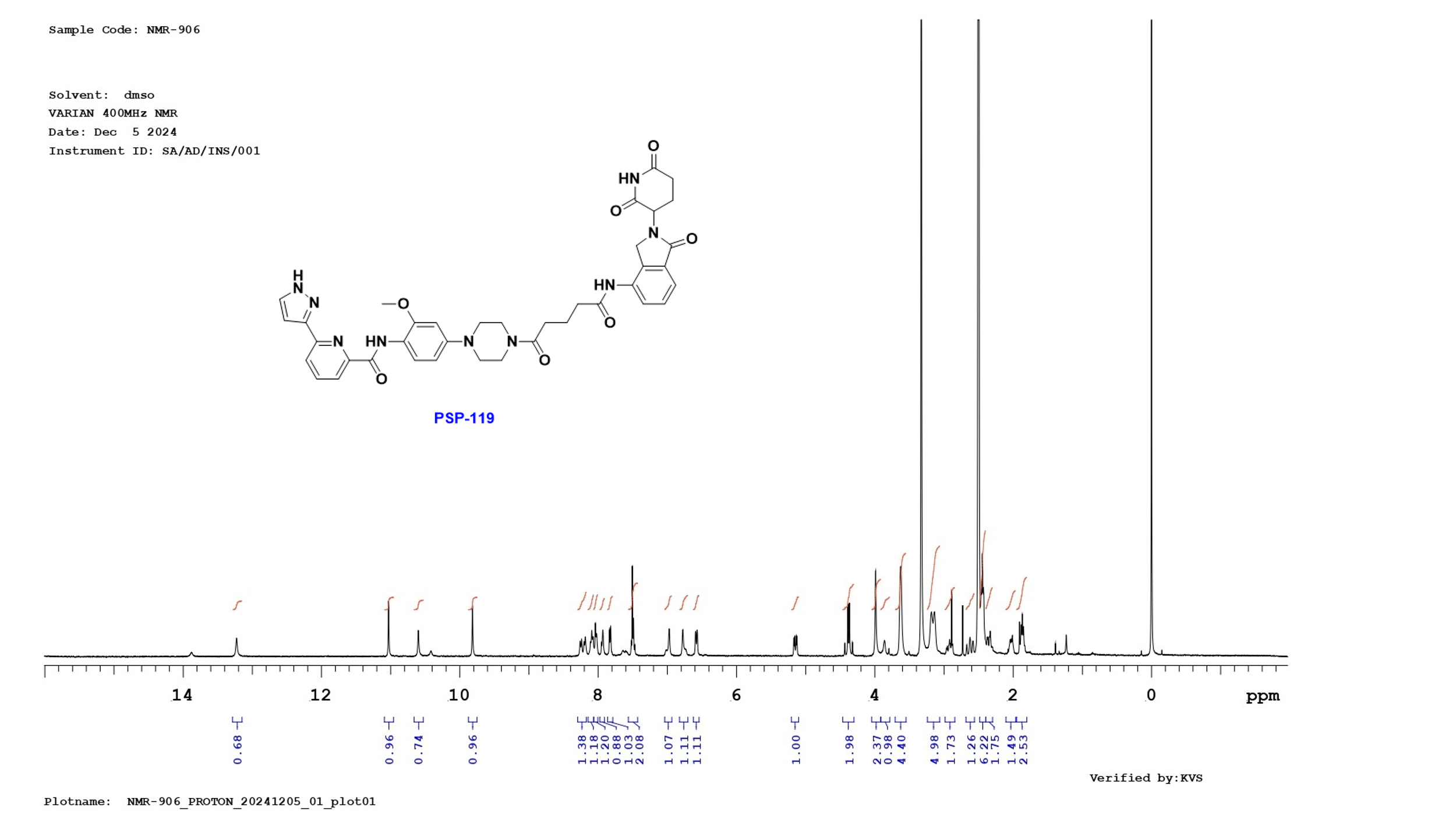

**Figure 9. MS spectrum of intermediate PSP-0119.**

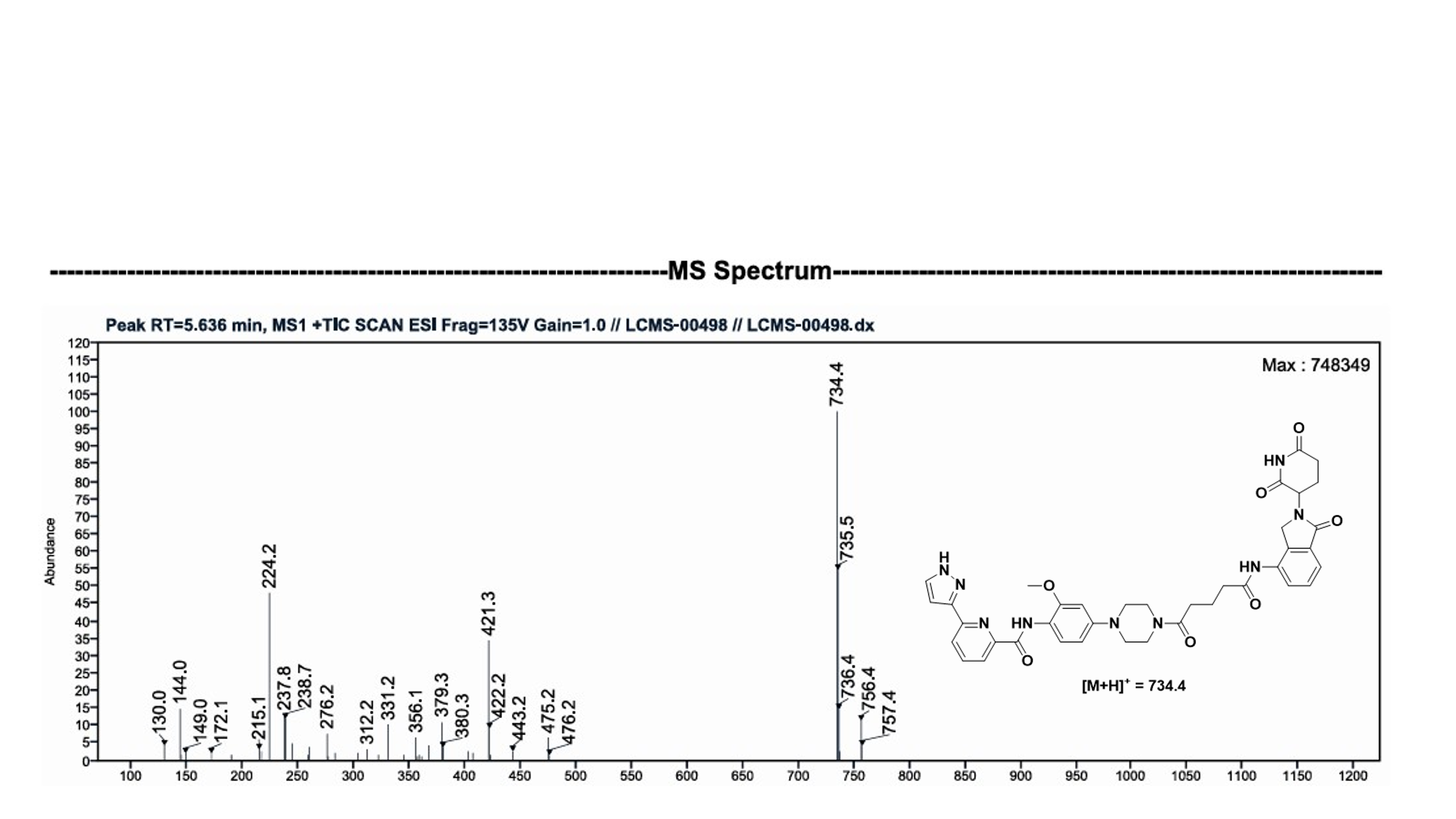

**HPLC of PSP-0119**

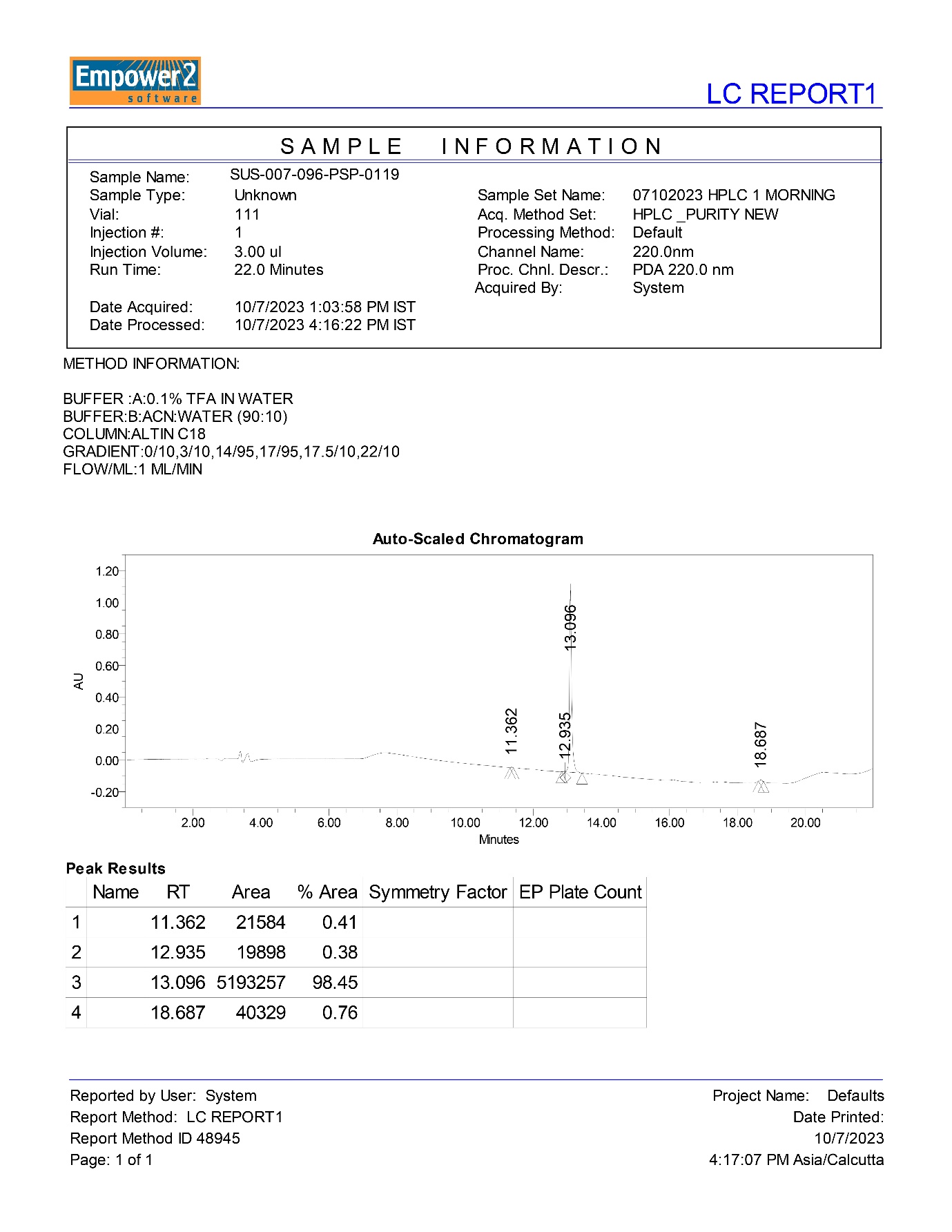

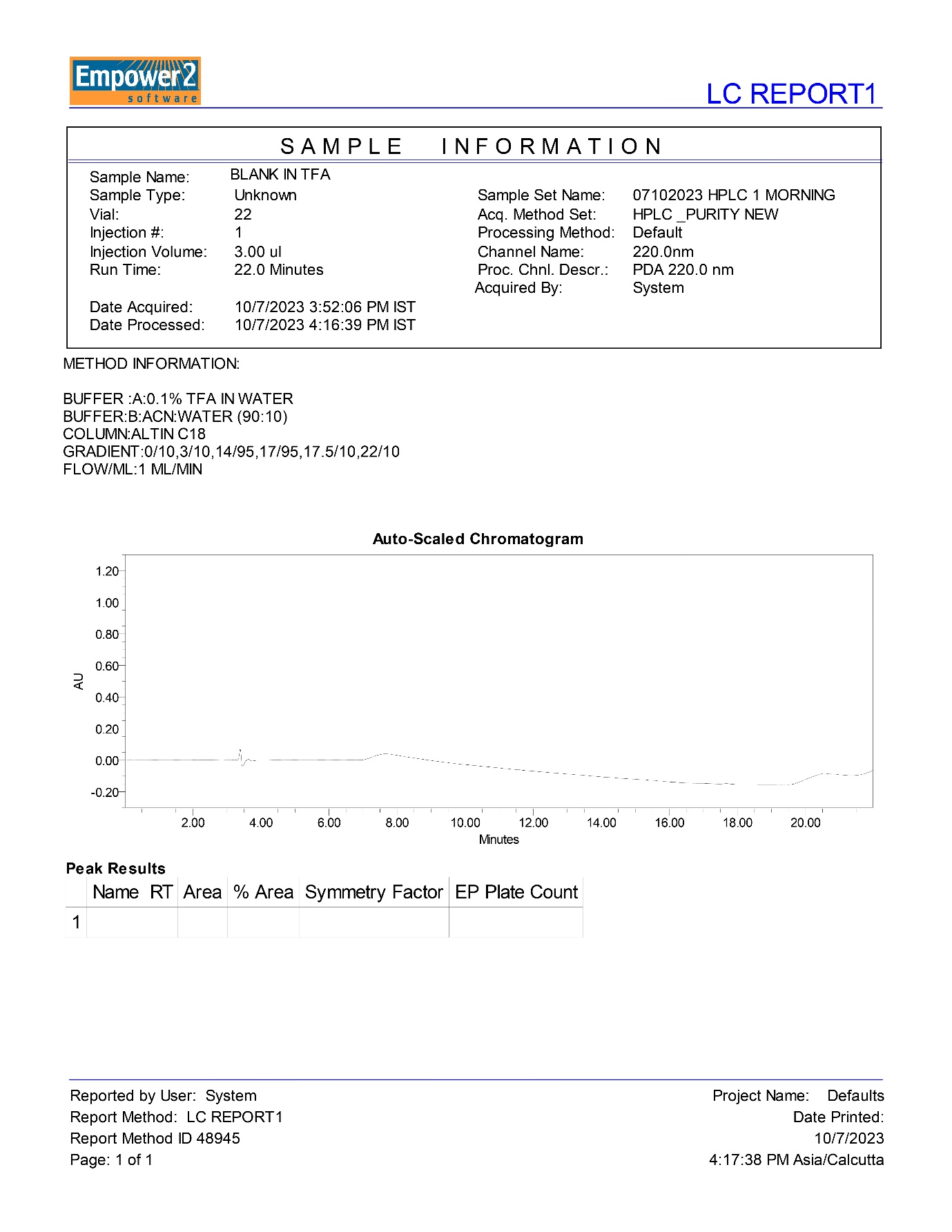

**Figure 10. ^1^H NMR spectrum of intermediate PSP-0102.**

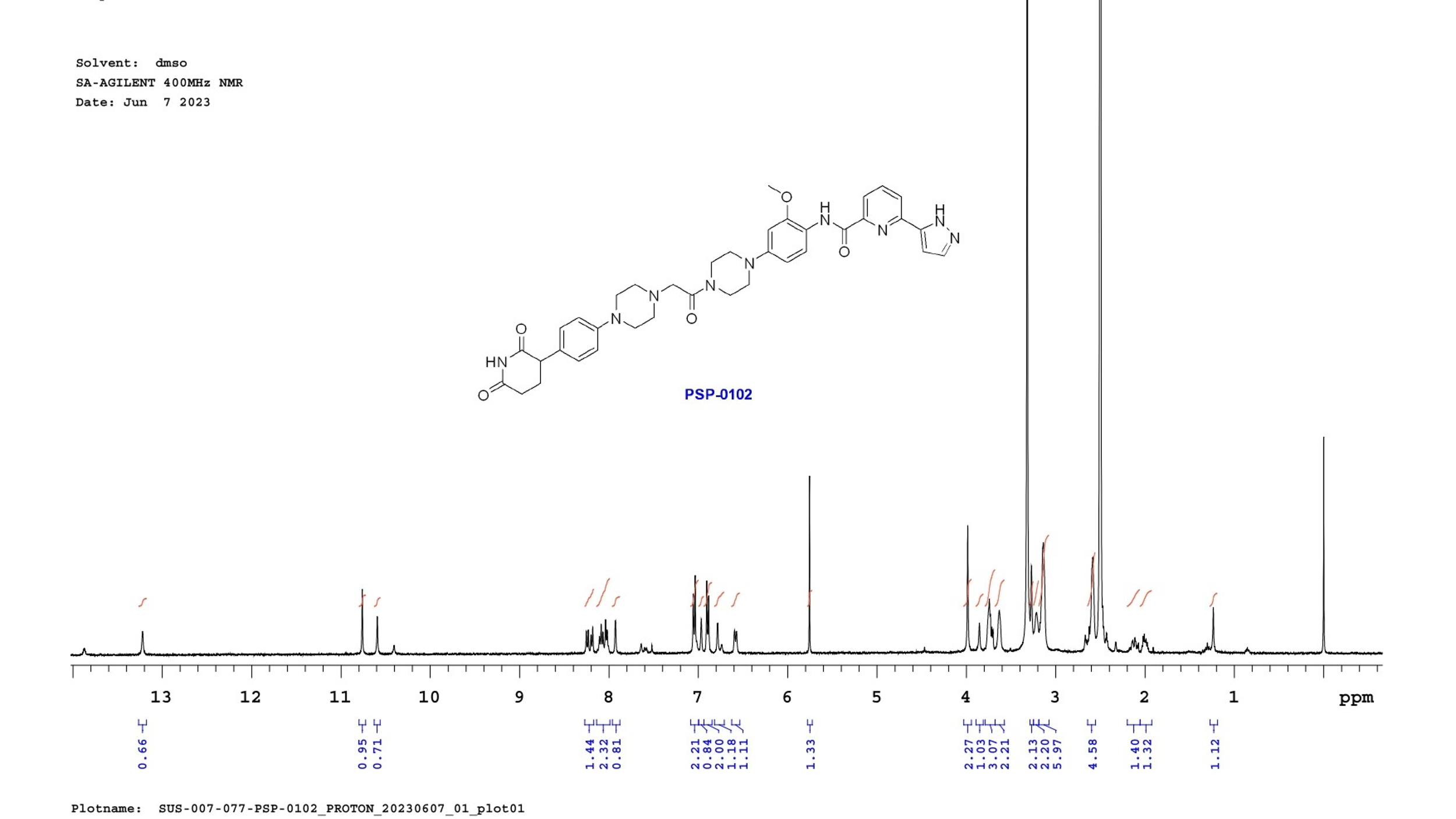

**Figure 11. MS spectrum of intermediate PSP-0102.**

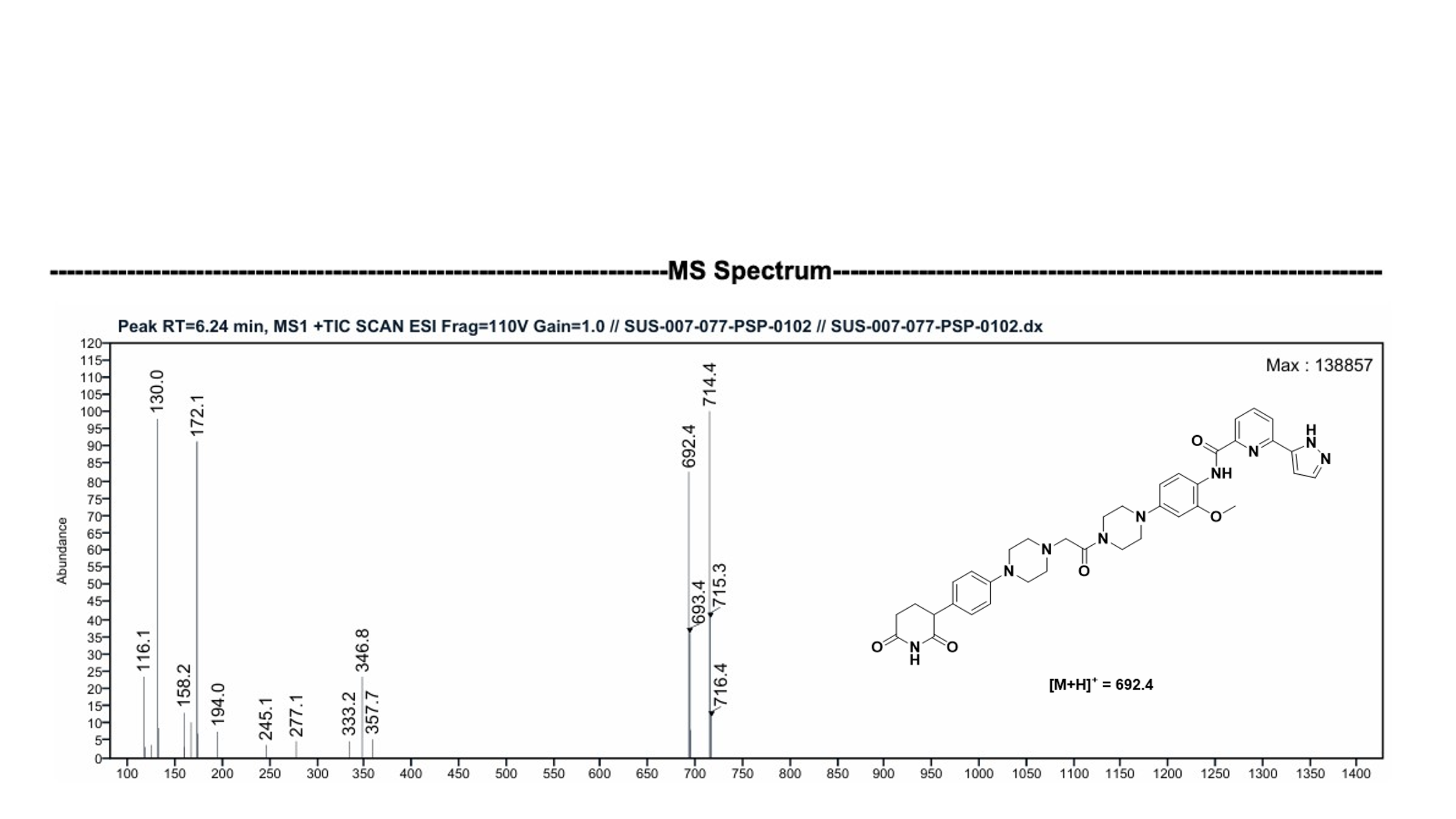

**HPLC data of PSP-0102**

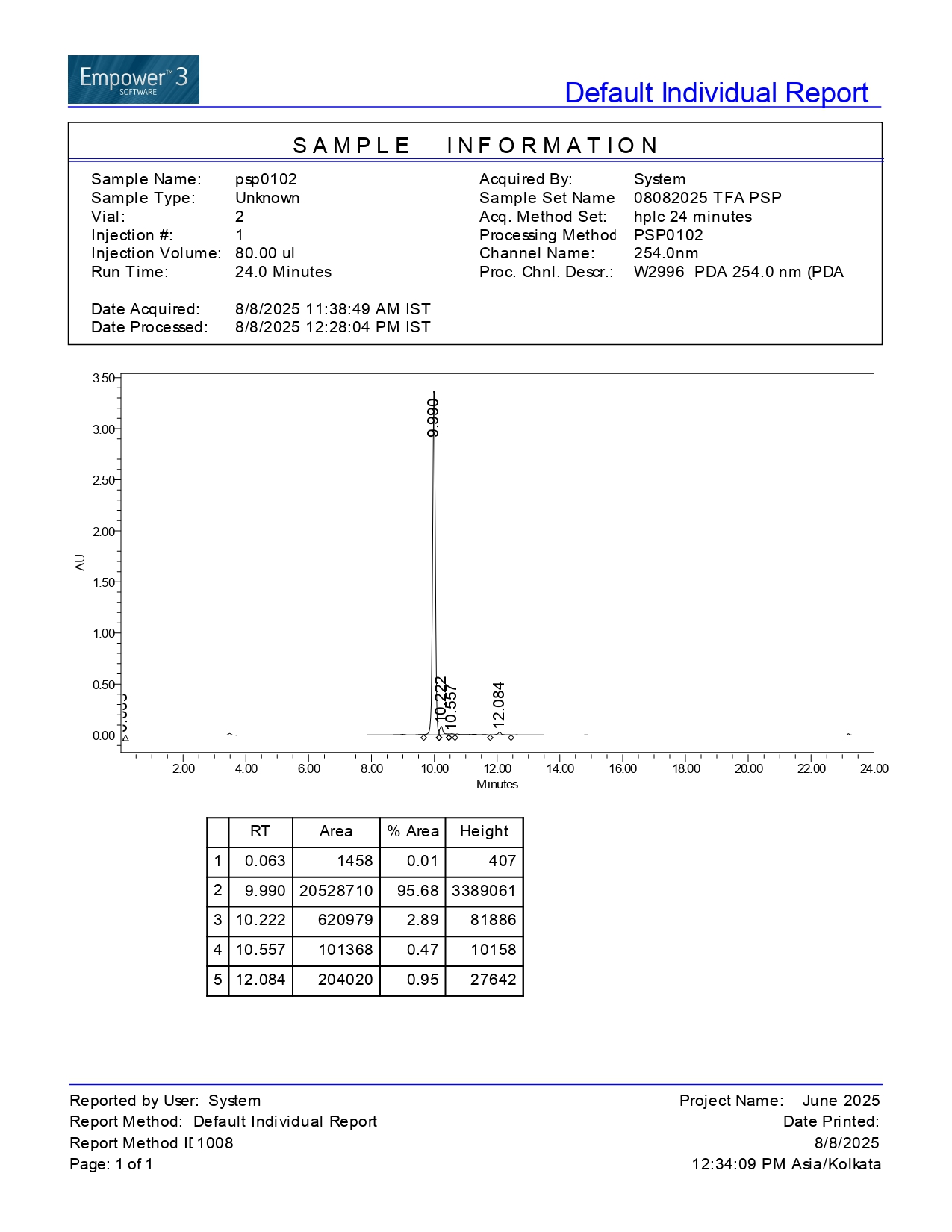

**Supplementary Figure-S2**

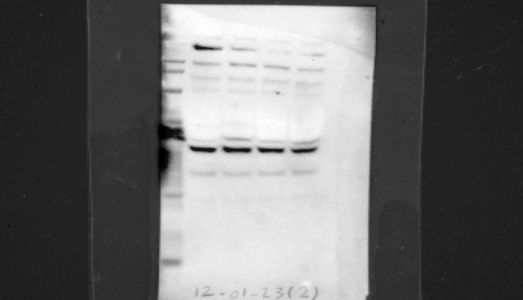

IRAK4-THP1

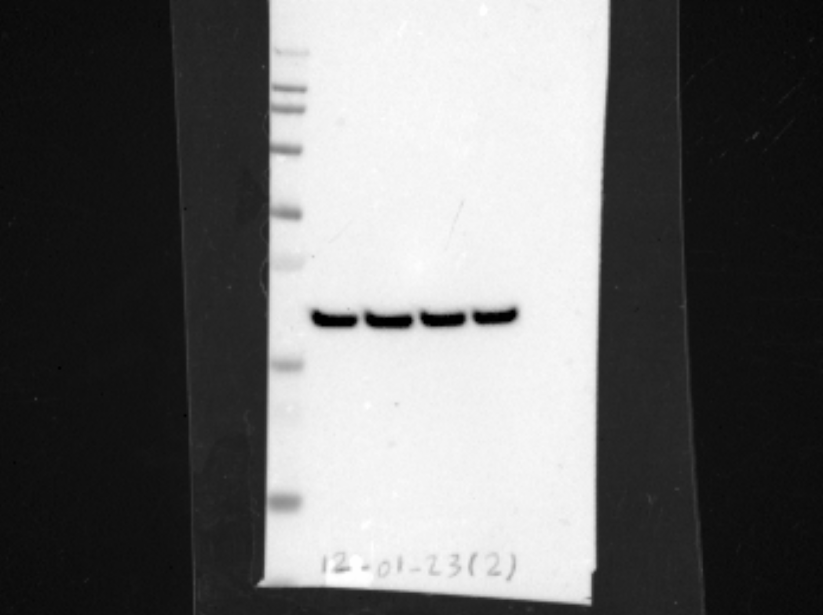

GAPDH

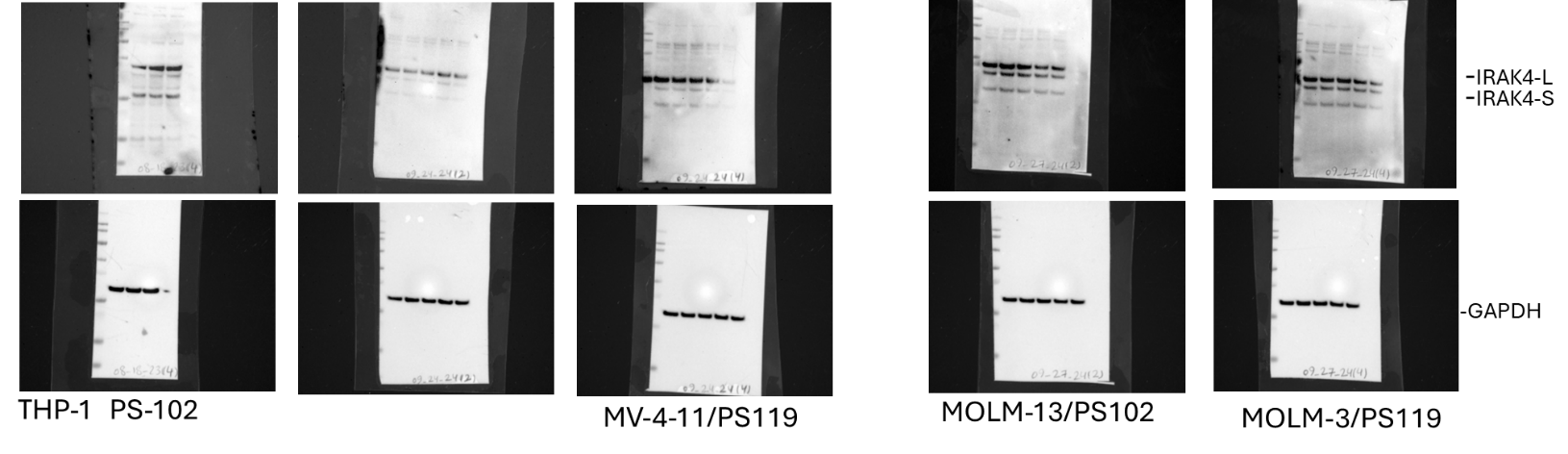

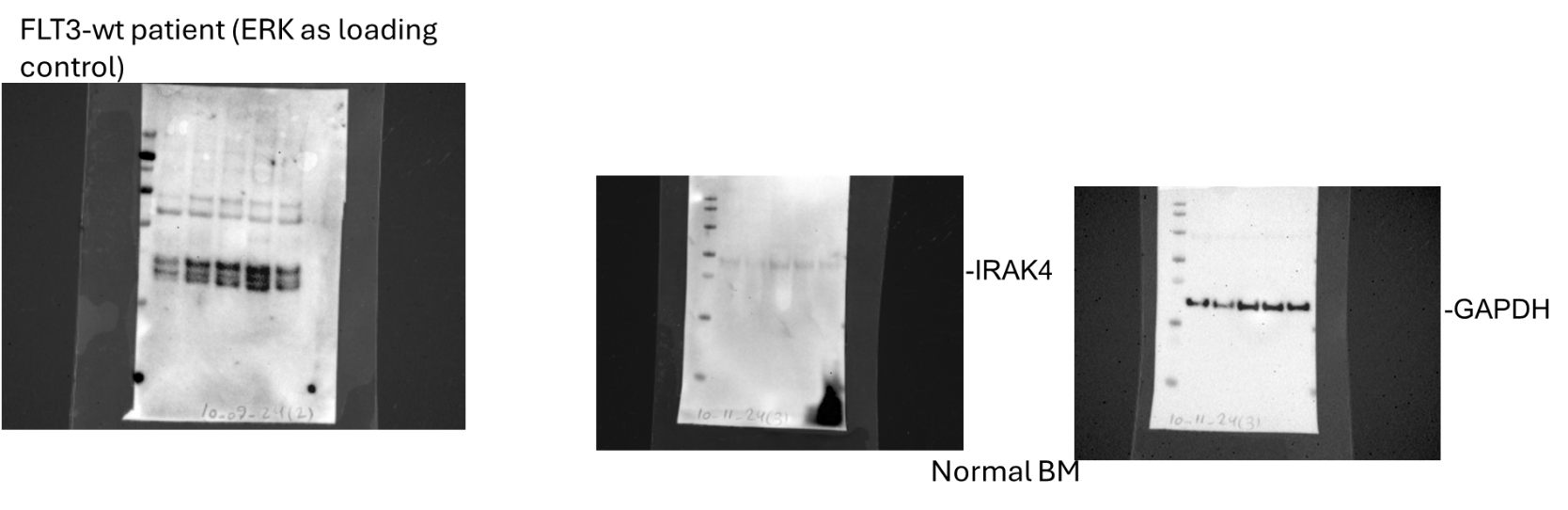

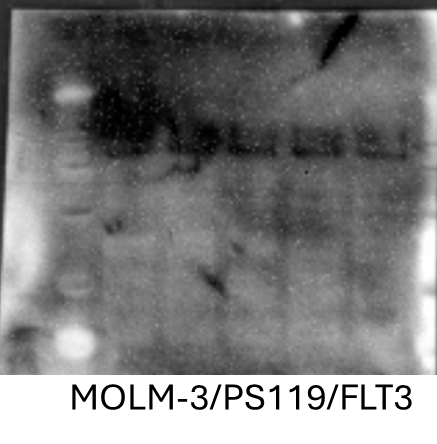

09-27-24(4) IRAK1

95

**Supplementary Figure-S3**

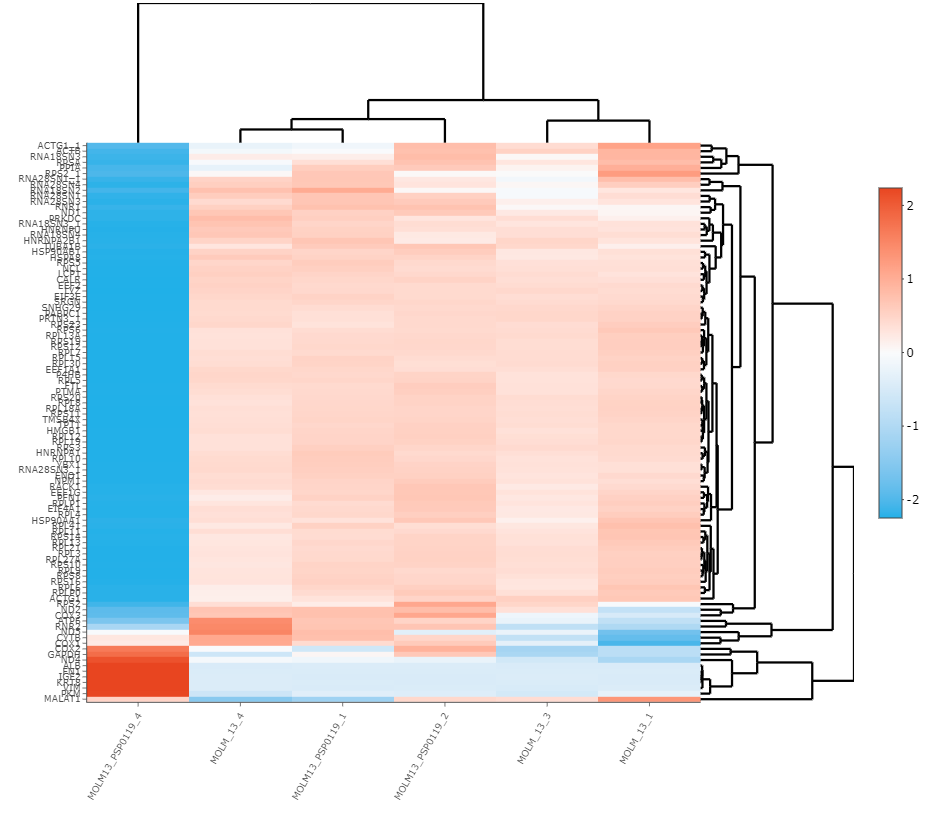

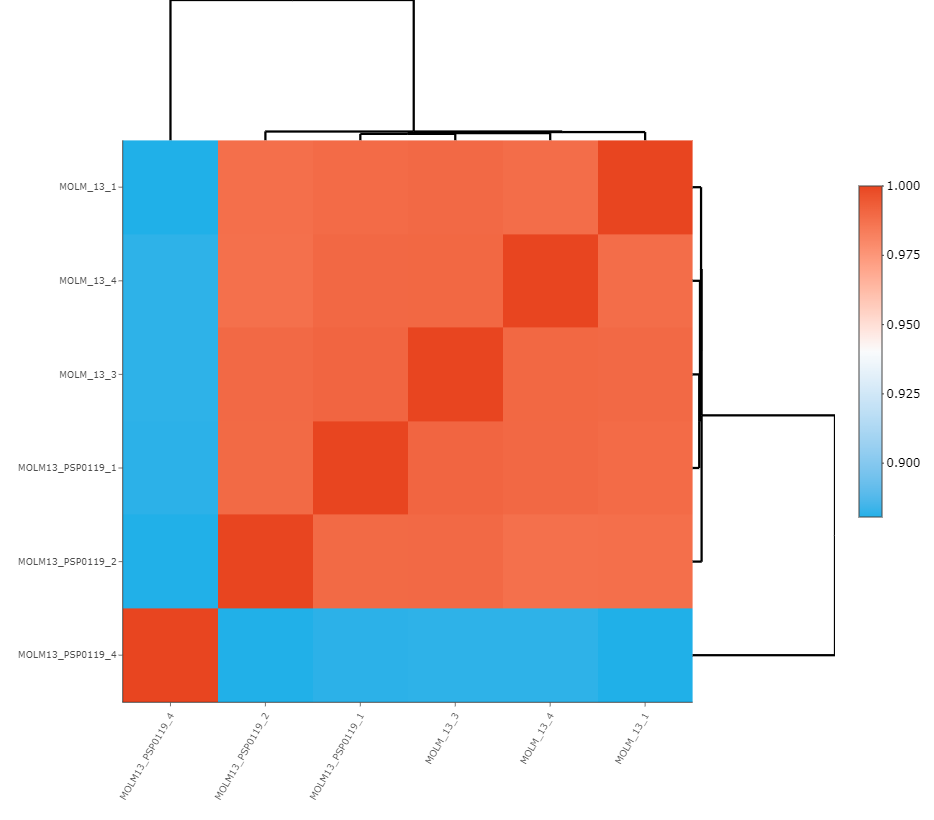

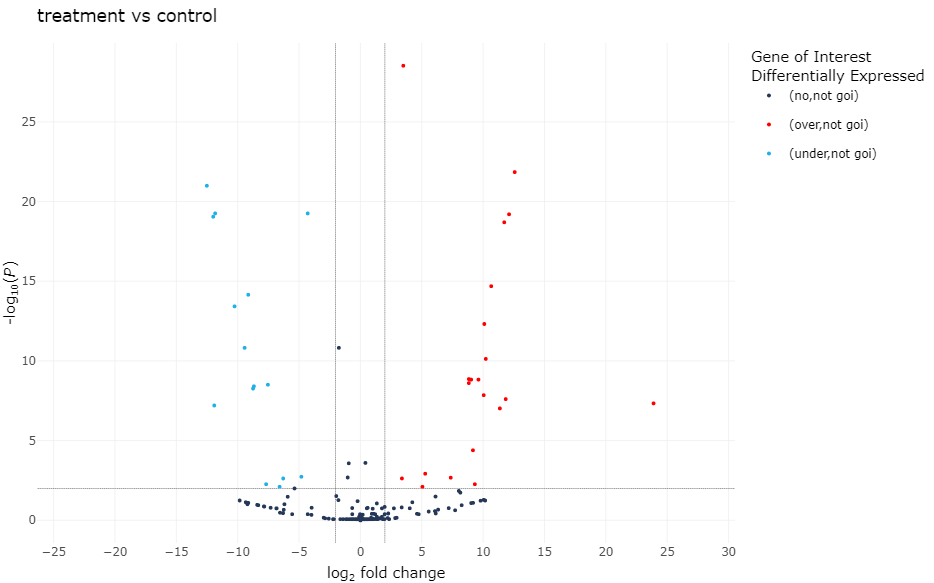

**Supplementary Figure- S4**

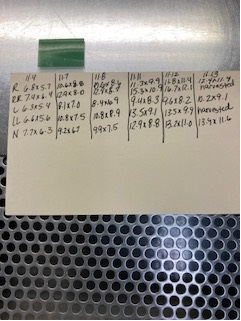

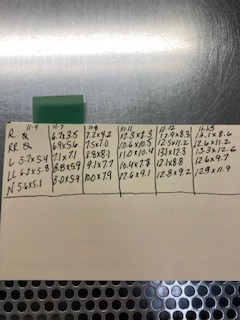

MOLm-13 xenograft in NSG mice

Treatment: PSP-0119 (10mg/kg, QD, IP) IRAK4 protac

vehicle

treatment
